# eIF5a reduction via decreased Klf5 leads to cell senescence by mitochondrial fission in VSMCs

**DOI:** 10.1101/787812

**Authors:** Dong Ma, Bin Zheng, He-liang Liu, Yong-bo Zhao, Xiao Liu, Xin-hua Zhang, Qiang Li, Wei-bo Shi, Toru Suzuki, Jin-kun Wen

## Abstract

Though dysregulation of mitochondrial dynamics has been linked to cellular senescence, which contributes to advanced age-related disorders, it is unclear how Klf5, an essential transcriptional factor of cardiovascular remodeling, mediates the link between mitochondrial dynamics and vascular smooth muscle cell (VSMC) senescence. Here we show that Klf5 downregulation in VSMCs is correlated with rupture of abdominal aortic aneurysm (AAA), an age-related vascular disease. Mice lacking Klf5 in VSMCs exacerbate vascular senescence and progression of Ang II-induced AAA by facilitating ROS formation. Klf5 knockdown enhances, while Klf5 overexpression suppresses mitochondrial fission. Mechanistically, Klf5 activates eIF5a transcription through binding to the promoter of eIF5a, which in turn preserves mitochondrial integrity by interacting with Mfn1. Accordingly, decreased expression of eIF5a elicited by Klf5 downregulation leads to mitochondrial fission and excessive ROS production. Inhibition of mitochondrial fission decreases ROS production and VSMC senescence. Our studies provide a potential therapeutic target for age-related vascular disorders.

## Introduction

Cellular senescence is an important contributor to aging and age-related diseases, and the accumulation of cellular senescence is a main feature of aged organisms (Hernandez-Segura *et al*, 2018). Cellular senescence is traditionally defined as permanent cell cycle arrest in response to different damaging stimuli (Hernandez-Segura *et al*, 2018; Muñoz-Espín *et al*, 2014). In the cardiovascular system, the senescence of vascular smooth muscle cells (VSMCs) may be induced by different stimuli, such as angiotensin II ((Ang II), oxidative stress, inflammation, and DNA damage. On the other hand, VSMC senescence may also results in the loss of arterial function, chronic vascular inflammation, mitochondrial dysfunction, and the development of age-related vascular disorders, such as atherosclerosis, abdominal aortic aneurysm (AAA), hypertension, and diabetes (Chi *et al*, 2019; Lacolley *et al*, 2017).

Krüppel-like factor 5 (Klf5) is a zinc-finger transcriptional factor that regulates various cellular processes including proliferation, differentiation, development and apoptosis (Suzuki *et al*, 2009). In VSMCs, Klf5 is regulated by Ang II signaling and is an essential regulator of cardiovascular remodeling (Shindo *et al*, 2002). Ang II is known not only to regulate blood pressure and electrolyte balance but also to be involved in mediation of cell proliferation and oxidative stress, thus contributing to premature senescence (Valcheva *et al*, 2014; Dzau, 2001). During cardiovascular remodeling, Klf5 activates the expression of cell cycle promoting-genes, such as cyclin D1, cyclin B1, and growth factors and their receptors, such as PDGF, VEGF and VEGF receptor (Liu *et al*, 2010; Suzuki *et al*, 2005), with a concomitant suppression of negative cell cycle control gene p21 (Zheng *et al*, 2011; He *et al*, 2009). Our recent studies demonstrate that Klf5 is highly expressed in both macrophages and VSMCs of human and experimental mouse AAA. Moreover, Klf5 expression progressively increases with the aortic diameter expansion during Ang II-infusion-induced AAA formation (Ma *et al*, 2017). Despite the fact that 1) advanced age is a major risk factor for AAAs (Nordon *et al*, 2011), 2) VSMC senescence correlates with increased level of reactive oxygen species (ROS) (Przybylska *et al*, 2016), 3) Klf5 participates in development and progression of AAAs (Ma *et al*, 2017), it remains to be clarified whether and how Klf5 mediates the mechanistic and functional link between ROS generation and VSMC senescence.

Mitochondria play important roles in regulating critical cellular, physiological, and pathophysiological processes (Tezze *et al*, 2017; Anderson *et al*, 2018). Alterations in mitochondrial dynamics, which is regulated mainly by the processes of fission and fusion, are implicated in various human diseases including cancer and neurologic and cardiovascular diseases (Anderson *et al*, 2018; Abuarab et al, 2017; Vásquez-Trincado *et al*, 2016). Previous studies have demonstrated that dysregulation of mitochondrial dynamics is a key feature of aging (Anderson *et al*, 2018). Moreover, mitochondria are a major source of ROS, and mitochondrial dysfunction may leads to aberrant ROS production (Murphy 2009). Specifically, elevated ROS levels impair vascular cell life-span through the onset of cellular senescence (Valcheva *et al*, 2014) and have been demonstrated in human cerebral aneurysm (Aoki *et al*, 2009). These results indicate that dysregulation of mitochondrial dynamics can drive VSMC senescence and excessive ROS production. Although cardiomyocyte Klf5 was recently identified as a regulator of cardiac metabolism by directly activating transcription of PPARα and regulating lipid metabolism (Drosatos *et al*, 2016), whether and how Klf5 regulates mitochondrial dynamics is currently unclear.

It is well known that the dynamin-like GTPases mitofusin 1 and 2 (Mfn1, Mfn2) regulate the mitochondrial fusion process, and dynamin-related protein 1 (Drp1) plays a central role in the regulation of mitochondrial fission (Zhao *et al*, 2011). Eukaryotic translation initiation factor 5a (eIF5a) is localized not only to the nucleus but also to the mitochondria (Miyake *et al*, 2015) and is involved in the regulation of redox homeostasis (Alcolea *et al*, 2014). However, it remains unclear whether eIF5a, together with mitochondrial dynamics-related proteins, participates in the regulation of mitochondrial dynamics. Therefore, it is very intriguing to investigate whether eIF5a mediates functional link between Klf5 and mitochondrial dynamics as well as to explore the relationship between Klf5-regulated mitochondrial dynamics and VSMC senescence.

## Results

### Downregulation of Klf5 expression in VSMCs is correlated with the progression and rupture of aortic aneurysm

Because Klf5 is well known to mediate Ang II-induced vascular remodeling by stimulating VSMC proliferation (Suzuki *et al*, 2009), we sought to know how medial VSMCs were lost in Ang II-induced AAA. Thus, we determined the expression of Klf5 in human unruptured and ruptured AAAs. The results showed that Klf5 expression in ruptured AAA (4 cases) was significantly lower than that in unruptured AAA (22 cases), even though its expression was higher than that in normal abdominal aorta (8 cases) (Fig 1A and 1B). Further, confocal microscopy images were obtained by immunofluorescence staining with VSMC marker (α-actin, SMA) and Klf5, and showed that Klf5 was co-localized with VSMC marker in unruptured AAA (Fig 1C). Moreover, Klf5-positive VSMCs were hardly observed in ruptured AAA, while the percentages of Klf5-positive VSMCs were significantly higher in unruptured AAA than in the normal aorta (10.3 ± 2.2% vs. 2.4 ± 0.89%, Fig 1C and 1D).

**Figure 1.**
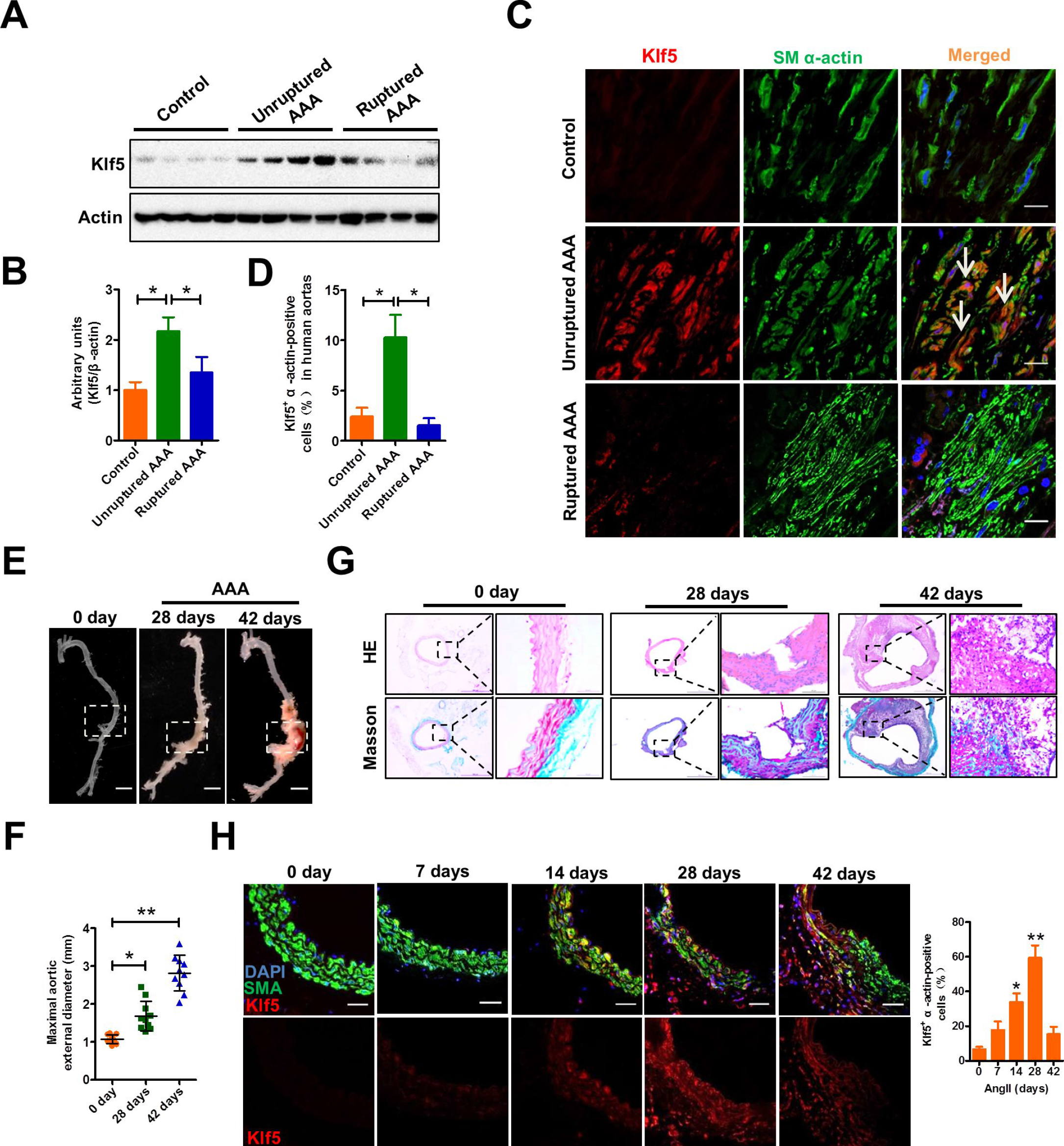
Klf5 expression is downregulated in ruptured human and experimental AAA. **A** Representative Western blots of Klf5 in human unruptured (n = 22) and ruptured (n = 4) AAA samples and in adjacent control aortas (control, n = 8). **B** Band intensities that were measured and normalized to β-actin. *P < 0.05 vs. control or unruptured AAA. **C** Immunofluorescence staining of smooth muscle (SM) α-actin, Klf5 and DAPI in human unruptured and ruptured AAAs and control. Arrows indicate Klf5^+^ SM α-actin-positive cells. Scale bars = 50 μm. **D** The percentage of Klf5^+^ SM α-actin-positive cells. *P < 0.05 vs. control or unruptured AAA. **E** Representative photographs of AAAs from ApoE^−/−^ mice infused with Ang II (1000 ng/kg/min) for 0, 28 and 42 days. Scale bars = 2.5 mm. **F** Statistical analysis of the maximal aortic external diameter. n = 10 for each group. Data represent the mean ± SD. *P < 0.05 and **P < 0.01 vs. 0 day. **G** Hematoxylin-eosin (top row) and Masson trichrome staining (bottom row) of abdominal vessels from ApoE^−/−^ mice treated as in (**E**). **H** Immunofluorescence staining of SM α-actin (SMA) and Klf5 in the injured aortas of ApoE^−/−^ mice exposed to Ang II for the indicated days. Scale bars = 50 μm. Right: Statistical analysis of Klf5^+^ SM α-actin-positive cells. *P < 0.05 and **P < 0.01 vs. 0 day.

In further studies, AAA models were generated in ApoE^−/−^ mice by chronic infusion of Ang II for 28 or 42 days, showing a typical aneurysmal phenotype or aneurysm rupture (Fig 1E). The aortic external diameter of Ang II-infused mice for 28 and 42 days increased obviously compared with that of the control mice (Fig 1F). Histological analysis with hematoxylin-eosin and Masson’s trichrome staining revealed an obvious collagen deposition in the medial layer of unruptured AAA (28 days) or collagen breakdown in ruptured AAA (42 days) (Fig 1G). We also determined the expression of Klf5 in VSMCs of AAA models and found that Klf5 expression and Klf5-positive VSMCs were significantly increased at 14 and 28 days after infusion with Ang II and subsequently started to decrease by 42 days (Fig 1H). These results imply that Klf5 appears to be detrimental in the early phase of AAA formation but is protective in late phase, and that Klf5 downregulation in the late phase of AAA is correlated with the progression and rupture of aortic aneurysm.

### Smooth muscle cell-specific knockout of Klf5 (smcKlf5^−/−^) exacerbates the progression of Ang II-induced AAA by facilitating reactive oxygen species (ROS) formation

To further investigate the VSMC-specific functions of Klf5 in the pathogenesis of AAA, we generated mice lacking Klf5 specifically in SMCs (Klf5^f/f^/Sm22^Cre/+^). Because advanced age is a known risk factor for AAA formation (Raffort *et al*, 2017), we used young (3-month-old) and old (18-month-old) Klf5^flox/flox^ mice (control, equivalent to wild-type) or smcKlf5^−/−^ mice to establish experimental AAA models by chronic Ang II infusion (Fig 2A). As shown in Appendix Fig S1A and 1B, aging led to an increase in SA-β-gal–positive staining regions in the aortas of WT mice, and the Klf5 loss in VSMCs further increased the areas of SA-β-gal–positive staining. The incidence of AAA in old mice had a significant increase compared with young mice regardless of Klf5 deletion in VSMCs, Klf5 deficiency resulted in a further increase in AAA incidence (Fig 2B). The aortic diameter of young smcKlf5^−/−^ mice had a >30% increase compared to young WT mice (1.73 ± 0.31 mm vs. 1.25 ± 0.28 mm, P < 0.05), but there was no difference in the survival rates between young WT and smcKlf5^−/−^ mice. In old mice, smcKlf5^−/−^ mice also had a >30% increase in the aortic diameter compared to WT mice (2.05 ± 0.38 mm vs.1.71 ± 0.33 mm, P < 0.05), and the survival rates were also significantly less in smcKlf5^−/−^ mice than in WT mice (75.0% vs. 89.7%, P < 0.05) (Fig 2C and 2D). These results suggest that aging increases Ang II-induced AAA formation in apoE^−/−^ mice and that Klf5 deficiency in VSMCs further exacerbates the progression of Ang II-induced AAA. Moreover, hematoxylin-eosin and Masson staining showed that Klf5 deficiency in VSMCs decreased collagen deposition and local thickness of media in old smcKlf5^−/−^ mice compared with those of old WT mice or young smcKlf5^−/−^ mice (Fig 2E and 2F). Notably, the extensive breakdown of collagen was observed in the ruptured aortas of old smcKlf5^−/−^ mice (Fig 2E).

**Figure 2.**
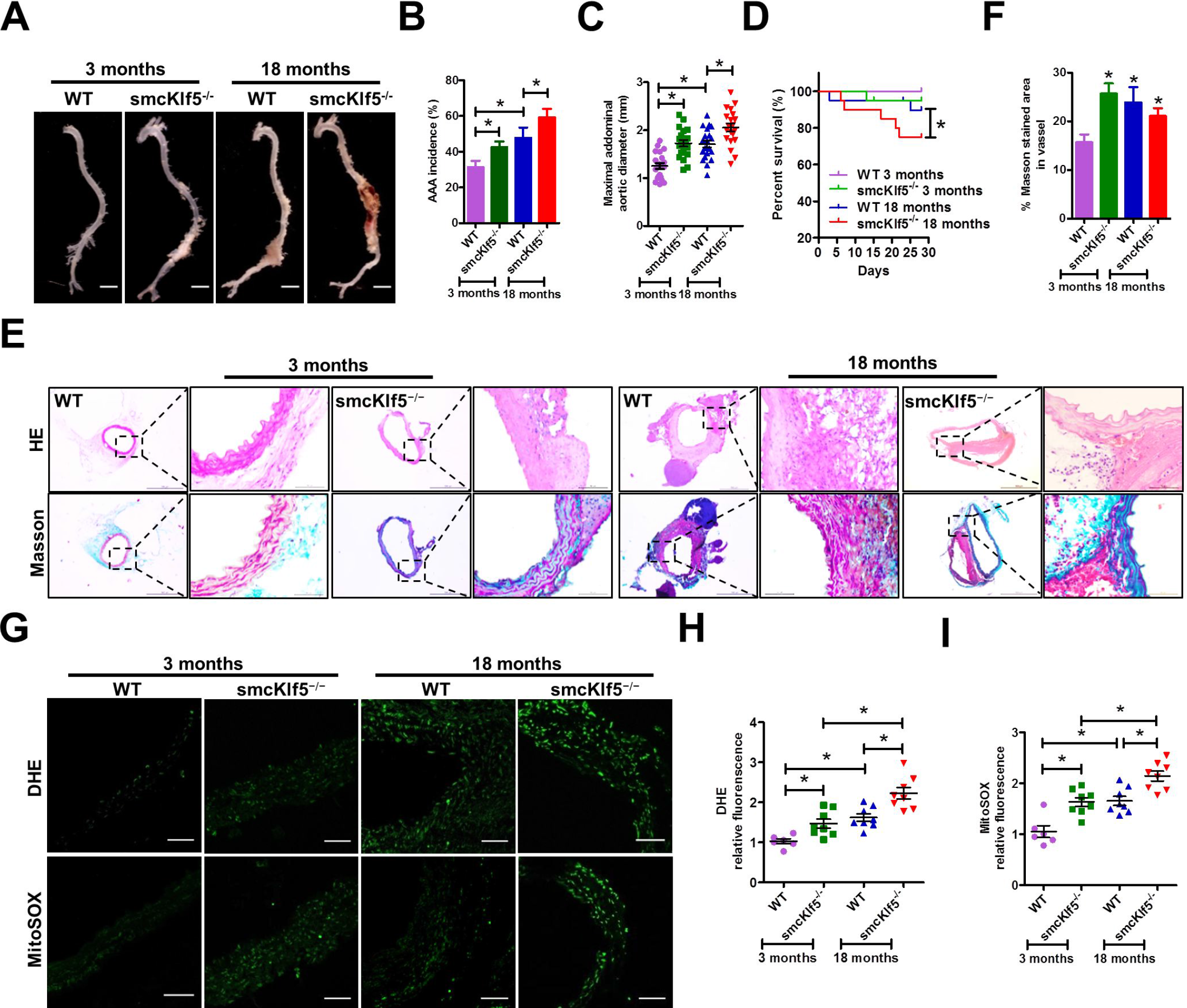
SMC-specific knockout of Klf5 (smcKlf5^−/−^) exacerbates the progression of AngII-induced AAA by facilitating ROS formation. **A** Representative photographs of AAAs induced by Ang II in young (3 months) or old (18 months) WT and smcKlf5^−/−^ mice. Scale bars = 2.5 mm. n = 20 for each group. **B-D** Statistical analysis of the incidence (**B**), the maximal aortic external diameter (**C**) and the survival rate (**D**). Data represent the mean ± SD. *P < 0.05 vs. young or old WT mice, log-rank test for **D**. **E** HE-(top row) and Masson-(bottom row) stained sections of the aortas from young or old WT and smcKlf5^−/−^ mice infused by Ang II for 28 days. Scale bars = 50 μm. **F** Quantification of Masson-stained area. n = 8~10. *P < 0.05 vs. young WT mice. **G** Dihydroethidium (DHE) staining (top) and mitoSOX staining (bottom) were performed to assess total cellular ROS and mitochondrial ROS. Scale bars = 50 μm. **H, I** Quantification of intracellular ROS levels based on measurement of fluorescence intensity. Data represent the mean ± SD. *P < 0.05 vs. WT or young smcKlf5^−/−^ mice, n = 8 for each group.

Because ROS plays a central role in the progression of AAA ^27^, we investigated whether loss of Klf5 in VSMCs can affect ROS production. Thus, young and old WT or smcKlf5^−/−^ mice were infused chronically with Ang II, and dihydroethidium (DHE) staining (to measure total superoxide) and mitoSOX staining (to measure mitochondrial superoxide) were performed. The results showed that the levels of total ROS and mitochondrial ROS (mtROS) were significantly higher in the aortas of old mice than in the aortas of young mice regardless of Klf5 deletion. Knockout of Klf5 in VSMCs further enhanced total ROS and the mtROS generation not only in young mice but also in old mice (Fig 2G–2I). In addition, TUNEL staining showed that alterations of cell apoptosis in the aortas of different mice (Appendix Fig S2). These findings indicate that increased ROS production elicited by Klf5 ablation is correlated with the progression of Ang II-induced AAA.

### Klf5 downregulation leads to VSMC senescence

Ang II not only induces oxidative stress but also promotes VSMC senescence (Andrés 2014; Miao *et al*, 2017). The above results indicate that SMC-specific Klf5 ablation also enhanced ROS generation. We next investigated whether Klf5 is involved in Ang II-induced ROS production and VSMC senescence. Human VSMCs were consecutively treated with Ang II (100 nmol/L) for 0, 1, 3 and 5 days and then cellular senescence was examined by β-galactosidase (SA-β-gal) staining, a classical method for detecting cellular senescence. The results showed that SA-β-gal–positive cells were significantly increased when VSMCs were exposed to Ang II for 5 days (Fig 3A), indicating that chronic Ang II stimulation results in VSMC senescence. Because the main characteristic of senescence is cell proliferation arrest (Lakatta *et al*, 2003), we examined the expression of Ki67, a marker for proliferating cells. As a result, Ki67 levels were markedly increased 1 day after Ang II treatment but returned to the basal level by 3 days. On day 5, Ki67-positive cells were hardly observed in the cell culture (Fig 3B and Appendix Fig S3). These data demonstrate that Ang II stimulates proliferation when acted on VSMCs for a short time, but chronic stimulation of Ang II induces VSMC senescence. Further, we explored the relationship between Klf5 and Ang II-induced VSMC senescence and found that Klf5 expression was significantly induced 1 day after Ang II stimulation, and then began to drop on day 3. On day 5, Klf5 expression was hardly detected by Western blotting (Fig 3C), consistent with Ki67 expression changes. These findings suggest that there is a direct relationship between Klf5 downregulation and Ang II-induced SMC senescence.

**Figure 3.**
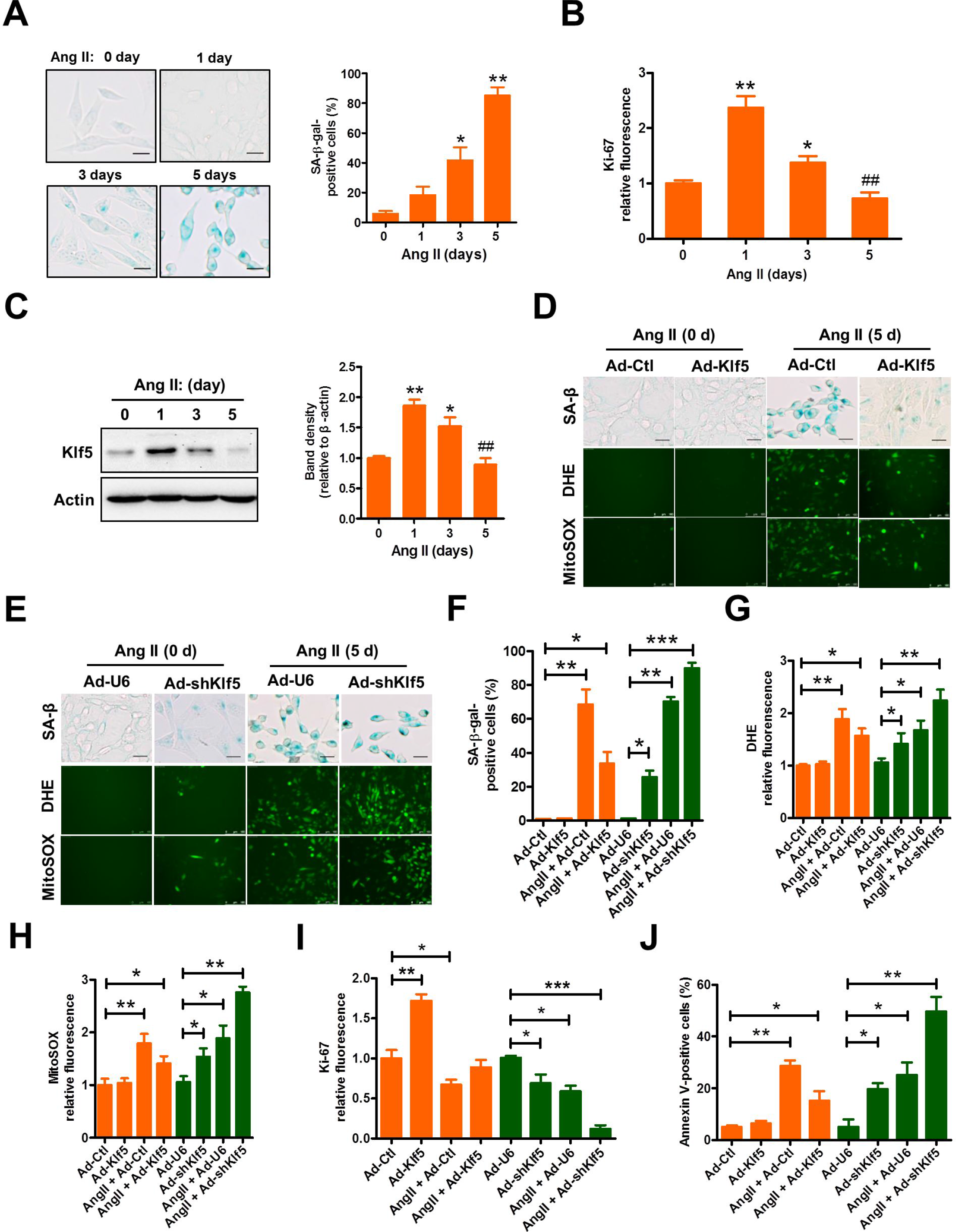
Klf5 downregulation leads to VSMC senescence. **A** Human VSMCs were stimulated with Ang II (100 nmol/L) for the indicated times. Senescence-associated β-galactosidase (SA-β-gal) staining (left) of Ang II-treated VSMCs and quantification β-gal-positive cells (right). Scale bars = 50 μm. Data represent the mean ± SD. *P < 0.05 and **P < 0.01 vs. 0 day (n = 3). **B** Quantification of Ki67-positive VSMCs treated as in (**A)**. Data represent the mean ± SD. *P < 0.05 and **P < 0.01 vs. 0 day, ^##^P < 0.01 vs. 1 day. **c** Representative Western blotting of Klf5 in VSMCs treated as in (**A).** β-actin serves as a loading control. Band intensities that were measured and normalized to β-actin were shown at the right side (n = 3). Data represent the mean ± SD. *P < 0.05 and **P < 0.01 vs. 0 day, ^##^P < 0.01 vs. 1 day. **D, E** SA-β-gal (top), DHE (middle) and mitoSOX staining (bottom) of VSMCs infected with the indicated constructs (**d**: Ad-Ctl and Ad-Klf5, **E**: Ad-U6 and Ad-shKlf5) and then treated with Ang II for 5 days. Scale bars=100 μm. **F-J** Quantification of SA-β-gal- (**F**), DHE- (**G**), mitoSOX (**H**)-, Ki67 (**I**) and Annexin V (**J**)-positive VSMCs. Data represent the mean ± SD. *p < 0.05, **p < 0.01, ***p < 0.001 vs. Ad-Ctl or Ad-U6.

In further experiments, we used a recombinant adenovirus (Ad-shKlf5 or Ad-Klf5) to knock down or overexpress Klf5 in human VSMCs (Appendix Fig S4). SA-β-gal staining for VSMCs showed that the number of SA-β-gal–positive cells was dramatically reduced in Klf5-overexpressing VSMCs treated with Ang II for 5 days compared with that of Ad-Ctl-infected cells, with a concomitant decline in the total ROS and mtROS (Fig 3D and Fig 3F–3H). Conversely, knockdown of Klf5 by Ad-shKlf5 significantly enhanced the number of aged VSMCs upon exposure to Ang II for 5 days, with a concomitant increase in the total ROS and mtROS (Fig 3E–3H). Correspondingly, Klf5 overexpression or knockdown significantly increased or attenuated the number of Ki67-positive cells in Ang II-treated VSMCs for 5 days (Fig 3I). Simultaneously, chronic stimulation of Ang II also induced apoptosis in Klf5-knocked down or overexpressed VSMCs, and Klf5 knockdown further increased the number of apoptotic cells (Fig 3J and Appendix Fig S5). Collectively, these results suggest that downregulation of Klf5 facilitates VSMC senescence induced by chronic stimulation of Ang II at least in part via increasing ROS generation.

### SMC-specific ablation of Klf5 leads to mitochondrial fission

To clarify the mechanism whereby ROS production was increased in Klf5-deficient VSMCs, we first identified the genes which are regulated by Klf5 in aortic tissues. To do this, whole transcriptome analysis was performed by RNA-seq of the aortic tissues from smcKlf5^−/−^ mice versus WT mice infused with Ang II for 28 days. As a result, 993 genes were found to be differentially expressed between smcKlf5^−/−^ mice and WT mice (≥ 2-fold change in expression level; P < 0.05) (Fig 4A). Gene ontology (GO) and KEGG pathway analysis revealed that downregulated genes by Klf5 deficiency in aortic VSMCs were strongly associated with ATP production, respiratory burst and hydrogen peroxide catabolic process (Fig 4B). To further validate these findings, we infected VSMCs with Ad-Klf5 or Ad-Ctl and used RNA-seq analysis to compare the gene expression profiles between Ad-Klf5- and Ad-Ctl-infected VSMCs. The results showed that a total of 504 upregulated genes (≥ 2-fold change in expression level; P < 0.05) were detected in Klf5-overexpressed VSMCs. On the basis of the GO and KEGG pathway analysis, the upregulated genes by Klf5, opposite to what was observed in the aortic tissues of smcKlf5^−/−^ mice, mainly involved the genes related to ATP biosynthetic process and cell redox homeostasis. Notably, mitochondrial dynamics-related genes, such as Drp1, Mfn1, Mtfr1, Fis1 and Pink1, were also significantly differentially expressed (Fig 4C). Using quantitative real-time PCR, we further validated the expression of mitochondrial dynamics-, redox homeostasis- and ATP biosynthesis-related genes and obtained the results similar to those seen in mRNA microarray analysis (Fig 4D). Importantly, we found that eIF5a, a regulator of cell redox homeostasis (Alcolea *et al*, 2014), was also highly upregulated in Klf5-overexpressed VSMCs, as shown by RNA-seq analysis and qRT-PCR (Fig 4C and 4D).

**Figure 4.**
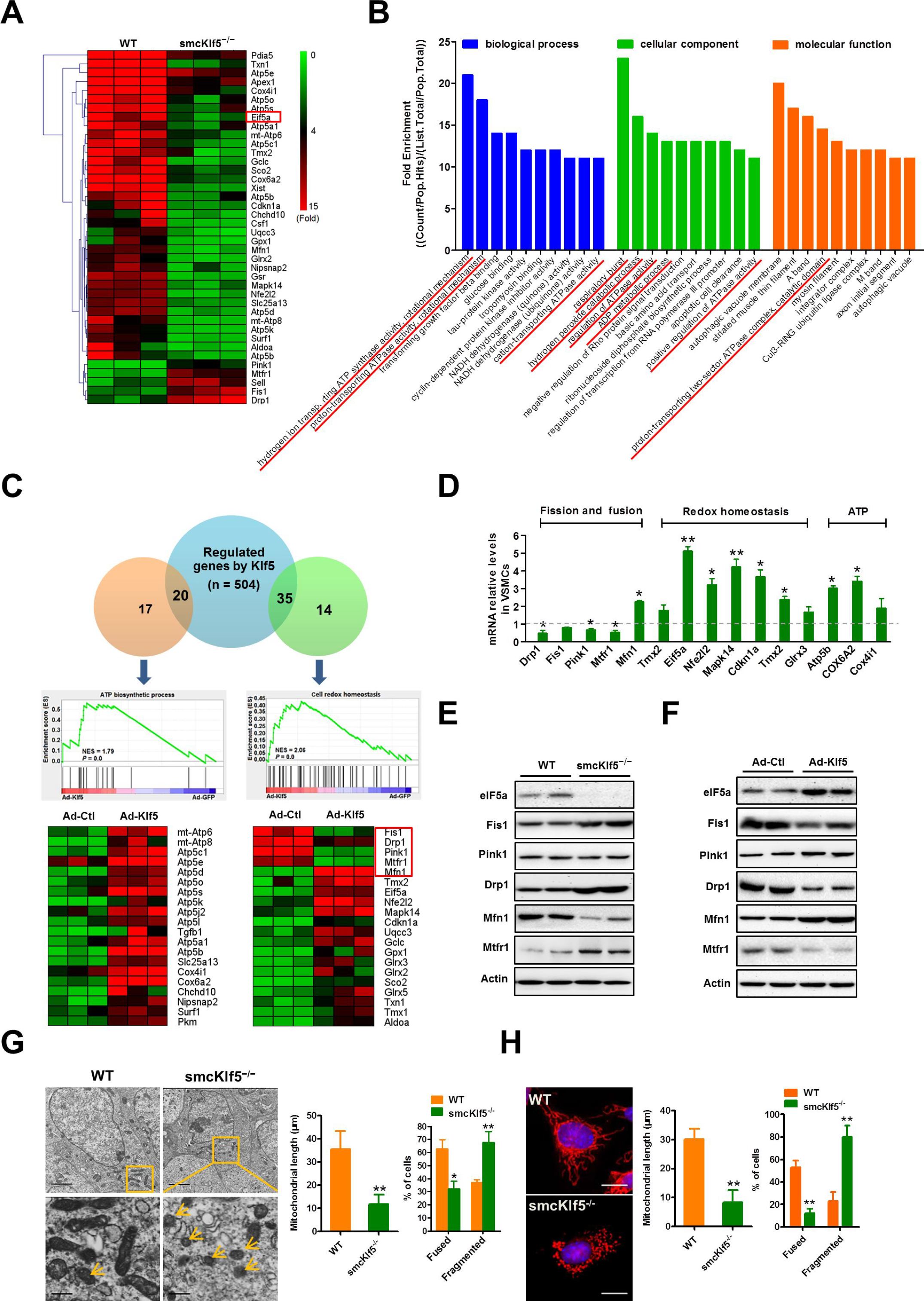
Klf5 deficiency in VSMCs leads to mitochondrial fission. **A**Hierarchical clustering of the top 40 differentially expressed genes identified by microarray analysis in AngII-injured aortas of WT and smcKlf5^−/−^ mice (n = 3). Red color indicates high relative expression and green color indicates low relative expression. **B** The Gene Ontology analysis for classification of the differentially expressed genes based on the biological process (blue), cellular component (green) and molecular function (red). **C** VSMCs were infected with Ad-Ctl or Ad-Klf5 for 36 h and then were analyzed by high-throughput mRNA sequencing. Gene Set Enrichment Analysis (GSEA) was used to identify the differential gene expression profiles between Klf5-overexpressing VSMCs and control VSMCs. **D** qRT-PCR validation of differential expression of a subset of genes including the fission and fusion-, redox homeostasis- and ATP biosynthesis-related genes. **E** Representative Western blot of eIF5a, Fis1, Pink1, Drp1, Mfn1 and Mtfr1 in the aorta of WT and smcKlf5^−/−^ mice, n = 4 for each group. **F** Representative Western blot of eIF5a, Fis1, Pink1, Drp1, Mfn1 and Mtfr1 in Ad-Ctl- and Ad-Klf5-infected VSMCs (n = 3). **G** Morphology of mitochondria in VSMCs of WT and Klf5^−/−^ mice was detected by transmission electron microscopy. Statistical data of the mitochondrial length and percentage of cells containing fused and fragmented mitochondria were shown in histogram at the right side. Data were expressed as mean ± SEM. n = 3, *P < 0.05 and **P < 0.01 vs. WT. **H** MitoTracker Red-stained mitochondria in WT and Klf5^−/−^ VSMCs. The nucleus was stained with DAPI. Right: the mitochondrial length and percentage of cells containing fused and fragmented mitochondria was quantified from more than 100 cells. Scale bars =10 μm. Data represent mean ± SEM, **P < 0.01 vs. WT.

Because the alteration of mitochondrial dynamics is associated with increased mitochondrial ROS production (Shenouda *et al*, 2011), we focused on the effect of Klf5 knockout or overexpression on the expression of eIF5a and mitochondrial dynamics-related genes. We found that mitochondrial fission factors Drp1, Fis1 and Mtfr1 were significantly upregulated, whereas fusion protein Mfn1 was downregulated in the aortic VSMCs of smcKlf5^−/−^ mice, without affecting the expression of several other proteins involved in the regulation of mitochondrial dynamics, as evidenced by Western blot analysis (Fig 4E). In contrast, overexpression of Klf5 in VSMCs produced the opposite effects on these gene expressions (Fig 4F). Because eIF5a is known to have a mitochondrial targeting property (Pereira *et al*, 2016) and its expression was dramatically decreased in the aortic VSMCs of smcKlf5^−/−^ mice (Fig 4A and 4E), we sought to know the effect of SMC-specific knockout of Klf5 on mitochondrial morphology. Thus, electron microscopy was used to examine the morphology of mitochondria. The results showed that the majority of mitochondria exist as rod-like shape (fused mitochondria) in the aortic VSMCs of WT mice. In contrast, in Klf5-deficient VSMCs, decreased mitochondrial length (fissed mitochondria) was observed, and the length and number of cells containing fragmented mitochondria were much more in Klf5-deficient VSMCs than in wild-type (WT) cells (Fig 4G). Also, mitochondrial morphology was visualized in WT and Klf5-deficient VSMCs by immunofluorescent staining of mitotracker. Similarly, the number of cells containing fused mitochondria was reduced, whereas fragmented mitochondria were increased in Klf5-deficient VSMCs (Fig 4H). Simultaneously, the ATP content in Klf5-deficient VSMCs also had a significant decrease compared with the WT cells (Appendix Fig S6). These findings clearly suggest that Klf5 deficiency in VSMCs leads to mitochondrial fission.

### Klf5 activates the transcription of eIF5a gene through direct binding to its promoter

Because eIF5a was downregulated in the aortic VSMCs of smcKlf5^−/−^ mice (Fig 4D and 4E), we sought to determine whether there exists a causal relationship between eIF5a downregulation and Klf5 deficiency as well as between eIF5a downregulation and time course of Ang II stimulation. First, we examined the effect of Ang II on the expression of eIF5a. In parallel with alterations in Klf5 expression (Fig 3C), the expression of eIF5a was markedly increased 1 day after Ang II treatment, but significantly decreased in Ang II-treated VSMCs for 3 and 5 days (Fig 5A). A similar result was obtained by qRT-PCR (Fig 5B). Further, we infected VSMCs with Ad-Klf5 or Ad-shKlf5 to overexpress or knock down Klf5. As expected, Klf5 overexpression increased, whereas Klf5 knockdown decreased eIF5a expression at both transcription and translation levels compared with their corresponding controls (Fig 5C and 5D). These results suggest that Klf5 plays an important role in the regulation of eIF5a expression.

**Figure 5.**
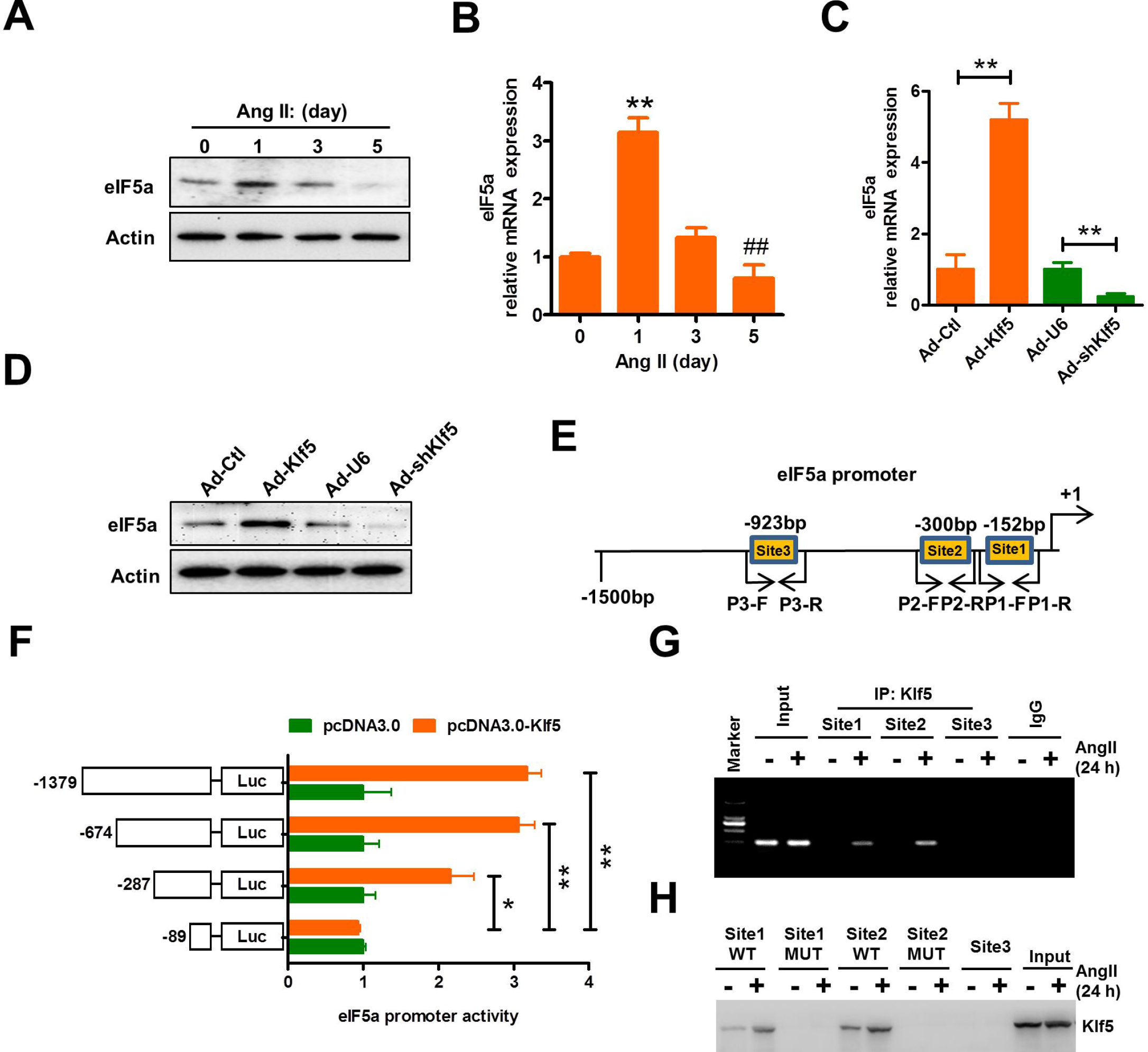
Klf5 activates the transcription of eIF5a gene through direct binding to its promoter. **A, B** VSMCs were stimulated with Ang II (100 nmol/L) for the indicated times (n = 3). The expression of eIF5a was analyzed by Western blotting (**A**) and qRT-PCR (**B**). β-actin was used as a loading control. **P < 0.01 vs. 0 day, ^##^P < 0.01 vs. 1 day. **C, D** VSMCs were infected with Ad-Ctl, Ad-Klf5, Ad-U6 and Ad-shKlf5 for 36 h. qRT-PCR (**C**) and Western blotting (**D**) detected mRNA and protein expression of eIF5a. β-actin was used as a loading control. **P < 0.01 vs. their corresponding controls. **E** A schematic map of the −1500 to +1 bp region of the human eIF5a promoter showing the position of 3 Klf5-binding sites and primers used for amplification. **F** 293A cells were transfected with the reporter directed by the eIF5a promoter containing different 5’-deletion fragments, and luciferase activity was measured. Data represent the relative eIF5a promoter activity normalized to pRL-TK activity. *p < 0.05 and **p < 0.01 vs. the reporter containing the −89 to +1 bp region. **G** VSMCs were incubated with or without Ang II (100 nmol/L) for 24 h. ChIP assay was then performed with antibody against Klf5. Nonimmune IgG was used as negative control. Immunoprecipitated DNA was amplified by PCR using the primers indicated as in (**E**). **H** An oligo pull-down assay was done with Ang II-treated VSMC lysates and biotinylated double-stranded oligonucleotide containing the Klf5-binding sites 1, 2 and 3 (WT and MUT) as probes. The DNA-bound protein was detected by Western blotting with anti-Klf5 antibody.

In further experiments, we investigated whether Klf5 regulates the expression of eIF5a by direct binding to its promoter. Using the JASPAR CORE database, we performed a Klf5-binding motif analysis on the eIF5a promoter, and identified three typical Klf5-binding sites in the −1500 to +1 bp of the 5’ upstream promoter of eIF5a gene (Fig 5E). Subsequently, a series of 5’-deletion mutants of the eIF5a promoter were constructed and tested by the luciferase activity assay. The results showed that enforced Klf5 expression markedly increased the activity of the eIF5a full-length promoter. Deletion of −674 to −287 bp or −287 to −89 bp of the eIF5a promoter region significantly decreased the activation by Klf5, indicating that Klf5-binding sites 1 and 2 between −300 bp and −152 bp are critical for the Klf5-mediated transcriptional activation of the eIF5a promoter (Fig 5F). To further validate these results, we carried out ChIP and oligonucleotide pulldown assays. The results showed that although the slight binding of Klf5 to the sites 1 and 2 of the eIF5a promoter could be detected under basal conditions, Ang II treatment obviously increased the recruitment of Klf5 to these two sites (Fig 5G and 5H). Mutation of these two sites abolished Klf5 binding (Fig 5H), indicating that the binding of Klf5 is specific. These results suggest that Klf5 activates the transcription of eIF5a gene by direct binding to its promoter and Klf5 downregulation is responsible for the decreased expression of eIF5a.

### eIF5a preserves mitochondrial integrity through interacting with Mfn1, downregulation of eIF5a elicited by Klf5 deficiency results in mitochondrial fission

Because alterations of mitochondrial dynamics, which is controlled by fission and fusion, induce the mtROS generation (Shi *et al*, 2018), and eIF5a is localized not only to the nucleus but also to the mitochondria (Miyake *et al*, 2015) and might be involved in the regulation of redox homeostasis (Alcolea *et al*, 2014), we wanted to determine whether eIF5a downregulation induced by Klf5 deficiency affects mitochondrial dynamics. Thus, the interaction of eIF5a with mitochondrial dynamics-related proteins, such as Mfn1, Drp1, Fis1 and Mtfr1, was examined in VSMCs stimulated chronically by Ang II. The co-immunoprecipitation assay confirmed that the interaction of eIF5a with fusion protein Mfn1was dramatically enhanced at 1 day after Ang II stimulation, but then markedly reduced at 5 days (Fig 6A). Also, an increased interaction between eIF5a and Mfn1 was detected 1 day after Ang II stimulation by *in situ* proximity ligation assay (Fig 6B). These changes are very similar to those of Klf5 and eIF5a expression following chronic Ang II stimulation. However, no significant interaction between eIF5a and fission factors Drp1, Fis1 or Mtfr1 was observed by co-immunoprecipitation assay (Appendix Fig S7). To provide additional confirmation that eIF5a interacts with Mfn1, co-immunofluorescence staining was performed using anti-eIF5a, anti-Mfn1 and mitotracker in eIF5a-overexpressed or knocked down VSMCs. Although eIF5a and Mfn1 co-localization in the mitochondria was detectable in empty vector-transduced VSMCs, eIF5a overexpression obviously increased their co-localization and facilitated the formation of network-like mitochondria (fused mitochondria), whereas eIF5a knockdown had the opposite effects (Fig 6C). These data suggest that eIF5a and Mfn1 are co-localized to the mitochondria and might play a functional role in the regulation of mitochondrial dynamics.

**Figure 6.**
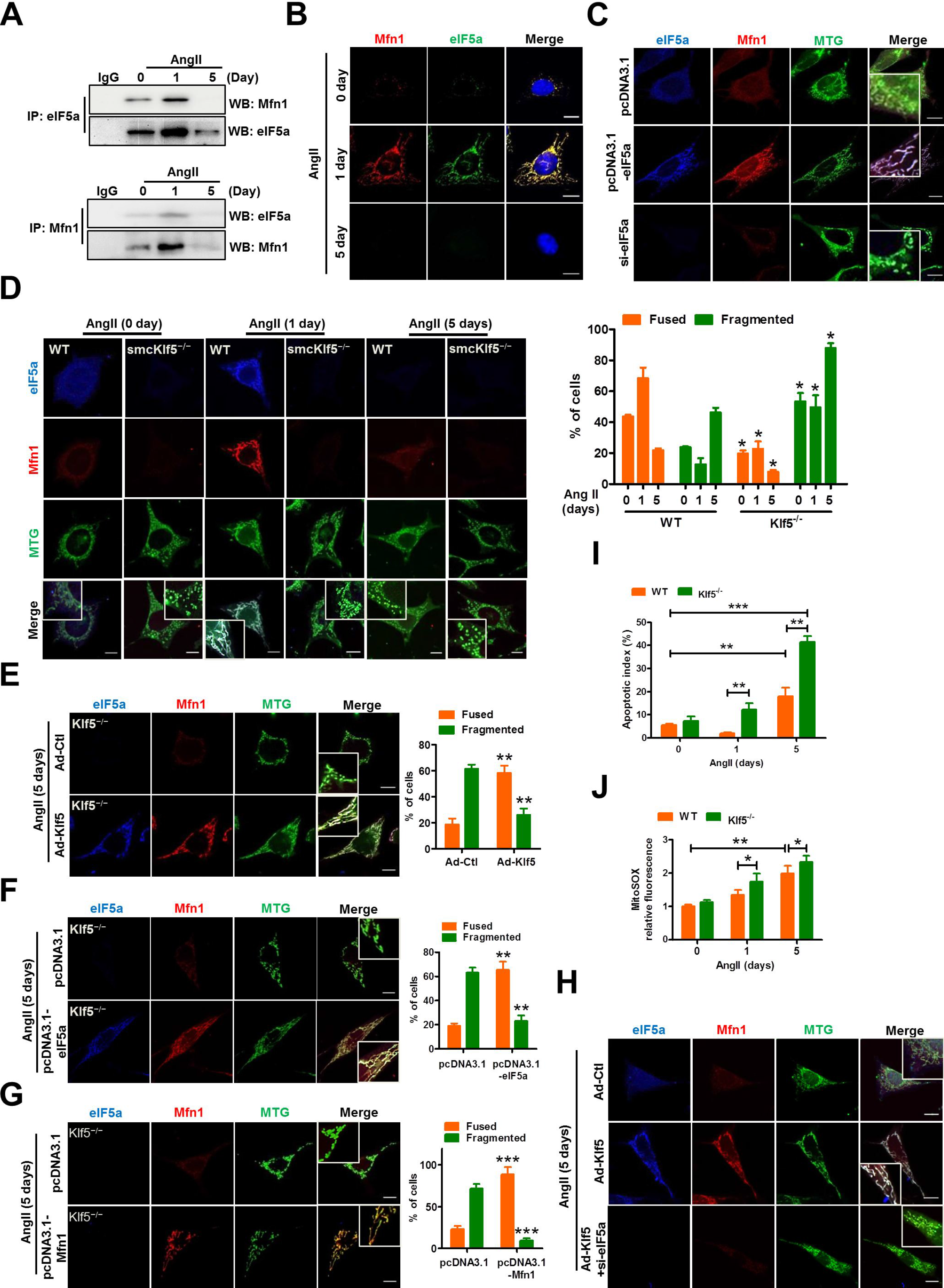
eIF5a interacts with Mfn1 and preserves mitochondrial integrity. **A** Reciprocal co-immunoprecipitation between eIF5a and Mfn1 in VSMCs treated with Ang II (100 nmol/L) for the indicated times. IgG was used as a negative control. **B** Duolink detection system (*in situ* PLA) was used to detect eIF5a and Mfn1 interaction *in situ* in VSMCs treated as in (**A)**. Scale bars = 10 μm. **C** VSMCs were transfected with pcDNA3.1, pcDNA3.1-eIF5a or si-eIF5a for 36 h. Colocalization of eIF5a with Mfn1 and MitoTracker Green (MTG) was detected by confocal microscopy. Scale bar = 10 μm. **D** WT and Klf5^−/−^ VSMCs were treated as described in (**A**). Fluorescence staining of eIF5a (anti-eIF5a, blue), Mfn1 (anti-Mfn1, red) and MTG was used to visualize mitochondrial morphology. Scale bars =10 μm. Percentage of cells containing fused and fragmented mitochondria is shown at the right side. Data represent mean ± SEM of 3 independent experiments in which 300 cells were analyzed. *P < 0.05 vs. WT. **E-G** Klf5^−/−^ VSMCs were infected with Ad-Ctl and Ad-Klf5 (**E**) or transfected with pcDNA3.1 and pcDNA3.1-eIF5a (**F**) or pcDNA3.1-Mfn1 (**G**) for 36 h. Mitochondrial morphology was visualized as described above. Scale bars = 10 μm. Right: Percentage of cells containing fused and fragmented mitochondria is shown as histograms. The data represent mean ± SEM of 3 independent experiments in which 300 cells were analyzed. **p <0.01, ***p < 0.001 vs. their corresponding control. **H** VSMCs were infected with Ad-Ctl, Ad-Klf5 or Ad-Klf5 plus si-eIF5a for 36 h and then treated with Ang II for 5 days. Mitochondrial morphology was visualized as described above. Scale bars = 10 μm. **I, J** Quantification of cellular apoptosis (**I**) and mitoSOX (**J**)-positive cells in WT and Klf5^−/−^ VSMCs treated as in (**D)**. Data represent mean ± SD, *p < 0.05, **p < 0.01, ***p < 0.001 vs. WT or 0 day.

Further, using cell immunofluorescent staining of mitotracker, eIF5a and Mfn1, we investigated the effects of chronic Ang II stimulation on mitochondrial dynamics in WT and Klf5-deficient VSMCs. When stimulated with Ang II for 1 day, the majority of mitochondria in WT VSMCs formed typical network-like structures. In contrast, when WT and Klf5-deficient VSMCs were treated with Ang II for 5 days, the majority of mitochondria displayed fragmented structures (fissed mitochondria) (Fig 6D). Correspondingly, the number of cells containing fused mitochondria was significantly increased in WT VSMCs treated with Ang II for 1 day. Conversely, upon exposure to Ang II for 5 days, the number of cells containing fragmented mitochondria was markedly elevated not only in WT cells but also in Klf5-deficient VSMCs (Fig 6D). These findings indicate that both loss of Klf5 and chronic Ang II stimulation lead to mitochondrial fission in VSMCs.

To corroborate these findings, we performed rescue experiments in which Klf5, eIF5a and Mfn1 were enforcedly expressed in Klf5-deficient VSMCs. The results indicated that not only Klf5-, but also eIF5a- or Mfn1-enforced expression could enable the fragmented mitochondria to fuse, forming a network-like structure (Fig 6E–6G). eIF5a knockdown could abrogate Klf5 overexpression-induced mitochondrial fusion (Fig 6H). Together, these data indicate that eIF5a or Mfn1 reside downstream of Klf5 and are involved in the regulation of mitochondrial dynamics. Additionally, we also found that knockdown of Mfn1 enhanced mitochondrial fission and abolished eIF5a enforced expression-promoted mitochondrial fusion (Appendix Fig 8A–8C). Moreover, increased mitochondrial fission was also accompanied by the increased apoptosis and the elevated mtROS levels in Ang II stimulated-VSMCs for 5 days (Fig 6I–6J).

### Inhibition of the mitochondrial fission decreases mitochondrial ROS production

To further clarify the relationship between mitochondrial fission and mtROS generation, we utilized two approaches to inhibit mitochondrial fission and then observed mtROS generation. First, compared with si-Ctl-transfected VSMCs, knockdown of Drp1, a key element in mitochondrial fission, by short interfering RNA (si-Drp1) could significantly suppress mitochondrial fission in VSMCs stimulated with Ang II for 5 days (Fig 7A). Further, we confirmed that pharmacological inhibition of Drp1 with Mdivi-1, a small molecule inhibitor of Drp1 (Bordt *et al*, 2017; Hoque *et al*, 2018) also markedly inhibited mitochondrial fission induced by chronic Ang II stimulation (Fig 7C). Correspondingly, the mtROS production was also reduced obviously in si-Drp1-transfected (Fig 7B) or Mdivi-1-treated VSMCs (Fig 7D).

**Figure 7.**
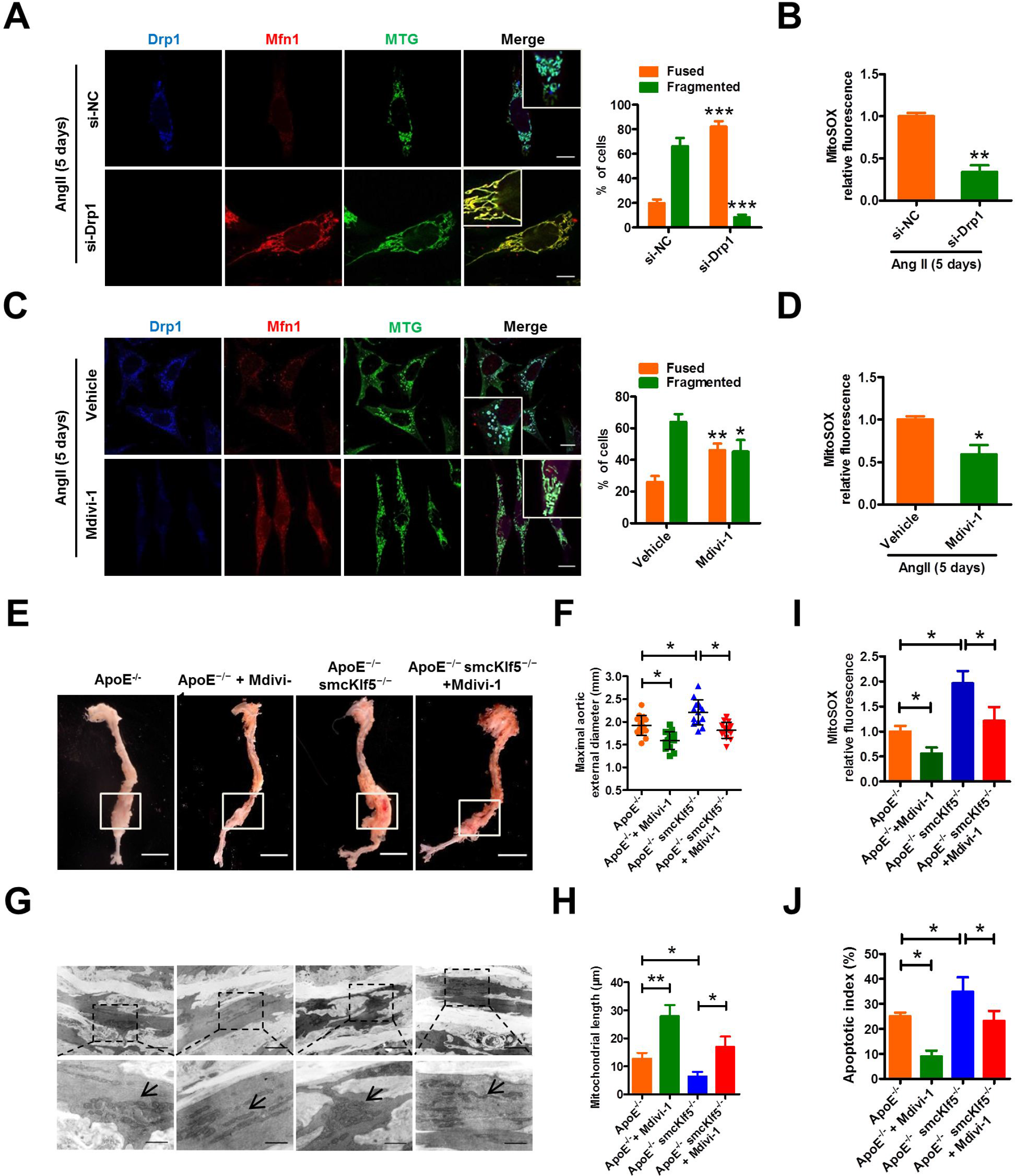
Inhibition of the mitochondrial fission decreases mtROS production. **A, C** VSMCs were transfected with si-NC and si-Drp1 (**A**) or treated with vehicle and Mdivi-1(inhibitor of Drp1) (**C**) for 24 h, and then stimulated with Ang II (100 nmol/L) for 5 days. Mitochondrial morphology was visualized by staining with MTG, anti-Drp1 and anti-Mfn1 antibodies. Scale bars = 10 μm. Right: Percentage of cells containing fused and fragmented mitochondria is shown as histograms. The data represent mean ± SEM of 3 independent experiments in which 300 cells were analyzed. *p < 0.05, **p < 0.01 and ***p < 0.001 vs. their corresponding control. **B, D** Quantification of mitoSOX-positive VSMCs treated with si-Drp1 (**B**) or Mdivi-1(**D**). Data are expressed as mean ± SEM. *p <0.05 and **p <0.01 vs. their corresponding control. **E** Representative photographs of AAAs induced by Ang II infusion for 28 days in ApoE^−/−^ and ApoE^−/−^ smcKlf5^−/−^ mice treated with or without Mdivi-1 for 4 weeks. Scale bars = 2.5 mm. **F** Statistical analysis of the maximal aortic external diameter. Data represent the mean ± SEM. *P < 0.05 vs. ApoE^−/−^ mice or ApoE^−/−^ smcKlf5^−/−^ mice. n = 8 for each group. **G** Transmission electron microscopy reveals mitochondrial morphology in AngII-injured aortas of ApoE^−/−^ or ApoE^−/−^ smcKlf5^−/−^ mice treated as in (**E**). **H** Statistical data of the quantified mitochondrial length in each group. Data are presented as the means ± SEM. *P < 0.05, **P < 0.01 vs. ApoE^−/−^ mice or ApoE^−/−^ smcKlf5^−/−^ mice. n = 3 in each group. **I**, **J** Graphical data show the mitoROS levels (**I**) and the percentage of apoptotic cells/total number of nucleated cells (**J**). All data are presented as the means ± SEM. *P < 0.05 vs. ApoE^−/−^ mice or ApoE^−/−^ smcKlf5^−/−^ mice.

**Figure 8.**
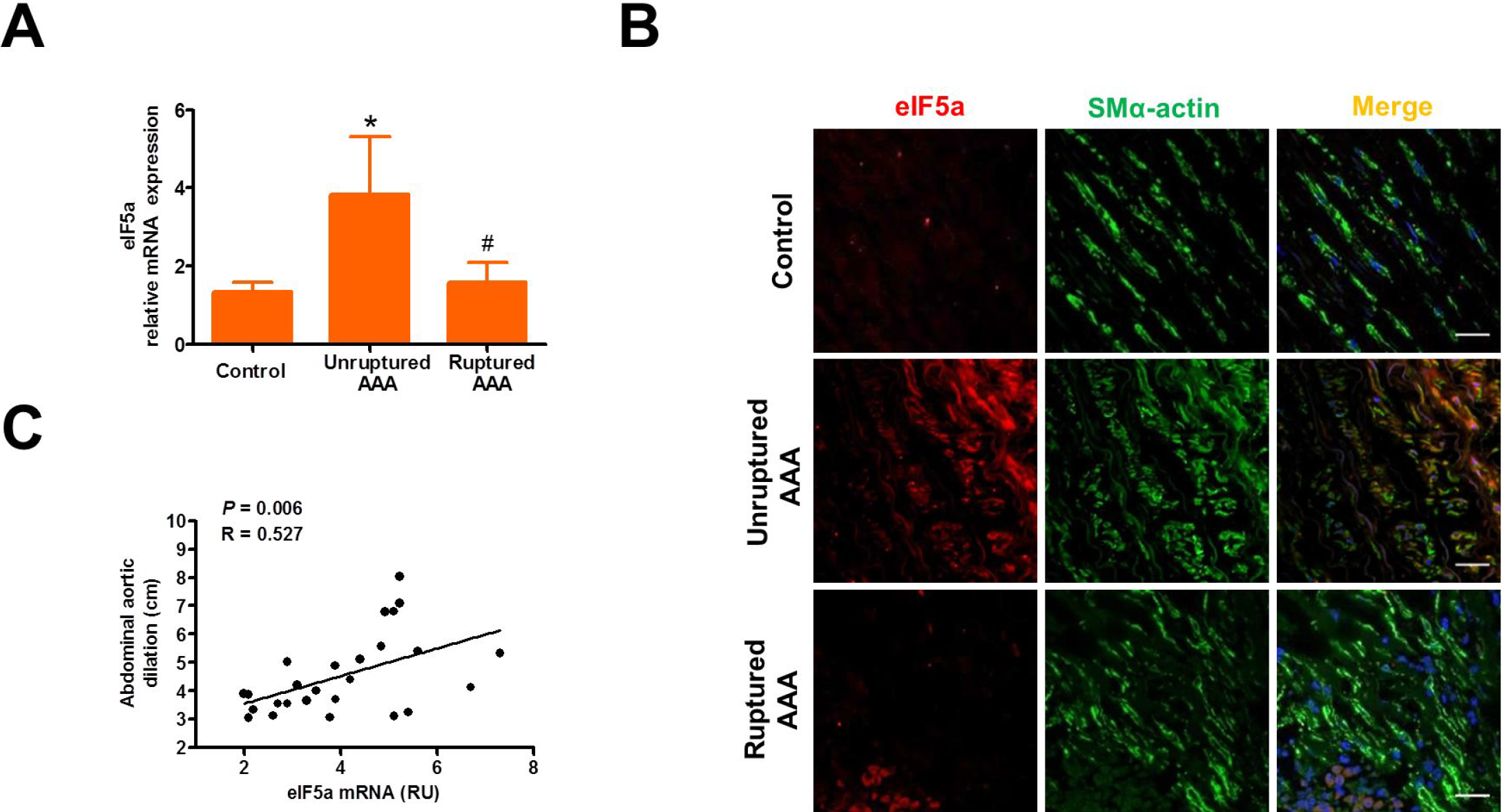
eIF5a is up-regulated in unruptured AAAs and decreased in ruptured AAAs. **A** qRT-PCR detected eIF5a mRNA in human unruptured (n = 22), ruptured (n = 4) AAA and in adjacent control aortas (control, n = 8), *P<0.05 vs. control, ^#^P < 0.05 vs. unruptured AAA. **B** Representative confocal fluorescence images of SM α-actin (green), eIF5a (red) and DAPI (blue) on human control, unruptured and ruptured AAA sections. Scale bars = 10 μm. **C** Correlation analysis between eIF5a mRNA level and AAA size. Spearman’s correlation coefficients were used to test the strength of a linear relationship between eIF5a mRNA and AAA size. Spearman’s correlation coefficient and P value are shown in graph. RU indicates relative unit.

To assess the effect of the inhibition of mitochondrial fission by Mdivi-1 on AAA formation, Mdivi-1 (25 mg/kg every other day) was administered by intraperitoneal injection to ApoE^−/−^ and ApoE^−/−^ smcKlf5^−/−^ mice, followed by chronic Ang II infusion for 28 days. The results showed that the aortic external diameter of ApoE^−/−^ smcKlf5^−/−^ mice had a > 20% dilation relative to ApoE^−/−^ mice. Importantly, administration of Mdivi-1 significantly decreased Ang II-induced aortic dilation regardless of Klf5 deletion in VSMCs (Fig 7E and 7F). Further, transmission electron microscopy was used to observe mitochondrial morphology and revealed that administration of Mdivi-1 to ApoE^−/−^ or ApoE^−/−^ smcKlf5^−/−^ mice significantly increased the mitochondrial length relative to their corresponding controls (Fig 7G and 7H). Also, we demonstrated the decreased mitochondrial fission was accompanied by the reduced mtROS production (Fig. 7i) and cell apoptosis (Fig 7J and Appendix Fig S9). Collectively, these results suggest that genetic and pharmacological *inhibition* of the mitochondrial fission can decrease mtROS production and AAA formation.

### eIF5a expression is up-regulated in unruptured AAAs and decreased in ruptured AAAs

To corroborate whether abnormal expression of eIF5a is involved in the development and progression of human AAAs, we examined the expression level of eIF5a in the normal aorta, unruptured AAA and ruptured AAA. The results showed that eIF5a expression was obviously higher in unruptured AAA than in the normal abdominal aorta, but its expression was significantly decreased in ruptured AAA compared with the unruptured AAA, as evidenced by qRT-PCR and immunofluorescence staining (Fig 8A and 8B). Moreover, eIF5a was localized to VSMCs in unruptured AAA tissues (Fig 8B). Statistical analysis revealed a strong correlation between the expression level of eIF5a and AAA size (Fig 8C). These results indicate that eIF5a downregulation induced by Klf5 deficiency in the aortic VSMCs is correlated with the progression and rupture of human aortic aneurysm.

## Discussion

The major findings of the present study were that *1*) Klf5 reduction in VSMCs is related to the rupture of human AAA; *2*) Loss of Klf5 in VSMCs exacerbates VSMC senescence by enhancing mitochondrial fission and ROS production; *3*) Klf5 activates the transcription of eIF5a gene through direct binding to the promoter of eIF5a, which in turn preserves mitochondrial integrity by interacting with Mfn1. eIF5a downregulation elicited by Klf5 deficiency results in mitochondrial fission; *4*) Inhibition of mitochondrial fission by genetic and pharmacological approaches decreases ROS production and VSMC senescence; and *5*) eIF5a expression is decreased in ruptured human AAA tissues.

Although Ang II is a key regulator of blood pressure and VSMC proliferation, chronic Ang II stimulation also induces senescence of vascular cells (Li *et al*, 2016; Kunieda *et al*, 2006). Moreover, Ang II is a potent mediator of oxidative stress and oxidant signaling leading to vascular premature senescence (Andrés, 2014). In this study, we showed that short-term exposure (1 day) of the cultured VSMCs to Ang II markedly increased Klf5 expression, while chronic Ang II stimulation (5 days) dramatically reduced Klf5 expression levels, accompanied simultaneously by an increase in the number of senescent VSMCs or a decrease in proliferating cells. In parallel with VSMC senescence, Klf5 knockdown or overexpression in VSMCs markedly enhanced or suppressed, respectively, Ang II-induced total ROS and the mtROS generation. It has been demonstrated that oxidative stress not only results in DNA damage, but also makes cells exit the cell cycle by inducing expression of negative cell cycle control genes p57kip2 and p21 (Valcheva *et al*, 2014; Zheng *et al*, 2011). Our results along with previous studies indicate that excessive ROS production induced by Klf5 deficiency leads to the lower proliferation capacity and the appearance of premature senescence in VSMCs. These are characteristic feature of the age-related vascular disorders, such as atherosclerosis, AAA, hypertension, and diabetes.

Mitochondria exist in a dynamic equilibrium between fragmented and fused states, and mitochondrial dynamics is controlled by fission and fusion (Anderson *et al*, 2018; Shi *et al*, 2018). Increased mitochondrial fission and/or attenuated fusion lead to mitochondrial fragmentation and disrupt cellular physiological function (Caja *et al*, 2017). Previous studies indicate that the increased mitochondrial fission leads to mitochondrial ROS (mtROS) generation and elevated apoptosis in endothelial cells (ECs) of diabetic patients (Shenouda *et al*, 2011; Yang *et al*, 2018; Galloway *et al*, 2012). Mitochondrial fragmentation, increased expression of the fission factor and reduced fusion protein levels were demonstrated in ECs of type 1 diabetic mice (Makino *et al*, 2010). In this study, we discovered that knockdown of Klf5 enhanced, while overexpression of Klf5 reduced mitochondrial fission induced by chronic Ang II stimulation. To further explore the relationship between mitochondrial fission triggered by Klf5 downregulation and ROS accumulation, we used microarray analysis to identify the mitochondrial dynamics-related genes regulated by Klf5. As expected, Drp1, Fis1 and Mtfr1 were significantly upregulated, whereas Mfn1 was decreased in Ang II-injured aortas of smcKlf5^−/−^ mice as well as in the aortas of older WT and smcKlf5^−/−^ mice, but the opposite results were obtained in Klf5-overexpressed VSMCs. These results suggest that loss of Klf5 in VSMCs leads to mitochondrial fission probably through affecting expression of mitochondrial dynamics-related genes. These are consistent with previous studies demonstrating that oxidative and other stresses frequently induce fission rather than fusion of mitochondria (Serasinghe *et al*, 2017). Accordingly, dysregulation of mitochondrial dynamics alters mitochondrial morphology and may affect mitochondrial function, accompanied by a decrease in ATP biosynthesis and an increase in mtROS generation.

Besides mitochondrial dynamics-related genes, eIF5a was highly expressed in Klf5-overexpressed VSMCs. Although eIF5a was originally believed to be a translation initiation factor that stimulates the initiation phase of protein synthesis by transient association with ribosomes (Benne *et al*, 1978), more recent studies show that eIF5a has numerous other functions. For example, hypusinated eIF5a modulates mitochondrial respiration and macrophage activation via promoting the efficient expression of a subset of mitochondrial proteins involved in the tricarboxylic acid cycle and oxidative phosphorylation (Puleston *et al*, 2019). eIF5a is also required for efficient translation of mRNAs encoding proteins with poly(Pro) tracts (such as PPP or PPG) by preventing ribosomes from stalling at such sequences (Pereira *et al*, 2016; Nguyen *et al*, 2015). Tyrosine sulfated-eIF5a is secreted from cardiac myocytes in response to hypoxia/reoxygenation and mediates oxidative stress-induced apoptosis (Seko *et al*, 2015). Inhibition of nuclear export protein exportin 1 (XPO1) causes eIF5a accumulation in the mitochondria and leads to the induction of apoptosis (Miyake *et al*, 2015). Reduced eIF5a function affects cell growth and autophagy via ATG3 protein synthesis (Lubas *et al*, 2018; Patel *et al*, 2009). Moreover, eIF5a was identified to be a significantly differentially expressed protein involved in the regulation of redox homeostasis (Alcolea *et al*, 2014). In this study, we first demonstrated that Klf5 directly bound to the eIF5a promoter and activated its transcription. Both Klf5 downregulation and chronic stimulation of VSMCs by Ang II dramatically attenuated the expression of eIF5a. Also, eIF5a expression was significantly reduced in aortic VSMCs of smcKlf5^−/−^ mice. These data suggest that eIF5a downregulation is responsible for Ang II-induced VSMC senescence, consistent with previous studies showing that a reduction of eIF5a content is associated with brain aging (Luchessi *et al*, 2008).

Importantly, using co-immunoprecipitation assay, *in situ* proximity ligation assay, and confocal immunofluorescence staining, we showed that eIF5a and the fusion protein Mfn1 were co-localized to the mitochondria. eIF5a overexpression obviously increased their co-localization and facilitated the formation of network-like mitochondria (fused mitochondria), whereas eIF5a knockdown had the opposite effects, accompanied simultaneously by a corresponding change in mitochondrial morphology. Together, these data suggest that eIF5a can be localized to the mitochondria and protects them from Klf5 deficiency-induced fragmentation. However, despite recent advances in our understanding of eIF5a functions, the excise mechanisms whereby mitochondrial integrity is maintained through interaction of eIF5a with Mfn1 need to be further elucidated.

Remarkably, genetic knockdown and pharmacologic inhibition of mitochondrial fission factor Drp1 could significantly attenuate mitochondrial fission and the mtROS production induced by chronic Ang II stimulation in VSMCs. Administration of Mdivi-1 to ApoE^−/−^ mice significantly decreased Ang II-induced AAA formation. These results are consistent with previous studies demonstrating that Mdivi-1 protects rat hippocampal neural stem cells against palmitate-induced oxidative stress and apoptosis by preserving mitochondrial integrity (Kim *et al*, 2017). These findings suggest that the prevention or inhibition of excessive mitochondrial fission may have therapeutic value in age-related vascular diseases.

Collectively, our studies discover a previously unrecognized role of eIF5a in the regulation of mitochondrial dynamics. Targeting the Klf5-eIF5a/Mfn1 regulatory pathway provides a potential therapeutic strategy for age-related vascular disorders.

## Materials and Methods

### Human tissue harvest

This study included 1) all patients with newly detected clinically intact (non-ruptured) aneurysms (n = 22), which underwent elective open AAA repair, and 2) those who were admitted for ruptured AAA (n = 4). AAA tissue specimens were obtained in the operating room from 26 male patients. Demographic and clinical characteristics of Patients with AAA were presented at Table 1. Nonaneurysmal infrarenal aortic wall tissue specimens were also obtained from organ donors to serve as nonaneurysmal controls (n = 8). Each of the surgical patients gave informed signed consent before donating tissue. Aneurysm size was measured on preoperative computed tomography angiograms. All tissue specimens were taken after a protocol approved by the Human Tissue Research Committee of Hebei Medical University.

**Table 1.** Demographic and clinical characteristics of Patients with abdominal aortic aneurysm (AAA)

| **Patients with AAA (n = 26 )** | **Data** |
| --- | --- |
| General characteristics |  |
| Age, years | 72 (65-82) |
| Sex male | 26 (100%) |
| Cardiovascular risk factors |  |
| Diabetes, n (%) | 7 (26.9%) |
| Arterial hypertension, n (%) | 19 (73.1%) |
| Dyslipidemia, n (%) | 16 (61.5%) |
| Smoking, n (%) | 14 (53.8%) |
| Clinical characteristics |  |
| Asymptomatic AAA, n (%) | 15 (57.7%) |
| Pain, n (%) | 7 (26.9%) |
| Ruptured AAA, n (%) | 4 (15.4%) |
| Imaging characteristics |  |
| AAA diameter, mm | 4.9 (3.1-8.0) |
| Comorbidities |  |
| Cardiac failure, n (%) | 5 (19.2%) |
| Renal failure, n (%) | 2 (7.7%) |
| Chronic obstructive pulmonary disease, n (%) | 6 (23.1%) |
| Preoperative full blood count |  |
| Red blood cells ($10^{12}/L$) | 4.4 (3.4-4.5) |
| Hemoglobin, g/dL | 12.7 (10.9-14.2) |
| Thrombocytes ($10^9/L$) | 214 (159-279) |
| Leukocytes ($10^9/L$) | 8.8 (7.3-10.8) |
| Neutrophils ($10^9/L$) | 5.8 (4.5-8.1) |
| Lymphocytes ($10^9/L$) | 1.8 (1.1-2.1) |
| NLR | 3.6 (2.4-5.8) |

Abbreviations: AAA, abdominal aortic aneurysm; NLR, neutrophil to lymphocyte ratio.

### Animal study

ApoE^−/−^ mice and Tgln-cre mice were purchased from Jackson Laboratory, and Klf5-flox mice were a gift from Huajing Wan (Wuhan University, China). We generated mice with smooth muscle cell-specific deletion of Klf5 by crossing Klf5-flox mice and Tgln-cre mice (smcKlf5^−/−^ mice) (Zheng *et al*, 2017). ApoE^−/−^ smcKlf5^−/−^ double-knockout mice were generated by crossing ApoE^−/−^ and smcKlf5^−/−^ mice. Homologous littermates with Klf5^+/+^ phenotype as WT controls were used for experiments. Genotyping was performed by PCR.

All male mice were housed and handled according to the guidelines of the local Animal Care and Use Committee at Hebei Medical University. To induce abdominal aortic aneurysms, we performed a mouse model of Ang II-induced AAA as previously described (Ma *et al*, 2017). Digital photographs of the abdomens were taken to measure the maximum external diameters. The abdominal arteries were harvested for analysis of RNA, morphology and histology.

Mitochondrial division inhibitor, Mdivi-1 was purchased from Enzo life sciences (Plymouth meeting, PA). Mdivi-1 was given as intraperitoneal injection in the dose of 30 mg per kg body weight every other day for 2 weeks. Since Mdivi was dissolved in di-methyl sulfoxide (DMSO), we gave similar dilution of DMSO in the normal saline as vehicle control. At the end of the treatment, animals were euthanized and aortas were harvested for analysis as above.

### Generation of primary cells and cell culture

Mouse primary VSMCs were isolated from aortas of 25 g male mice anesthetized intraperitoneally with urethane as described previously (Salmon *et al*, 2013). Mouse primary VSMCs were cultured in low-glucose Dulbecco’s modified Eagle’s medium (DMEM, Gibco Life Technologies, Rockville, MD) containing 100 units/ml of penicillin, 100 μg/ml of streptomycin and 10% fetal bovine serum (GEMINI, USA), in a humidified incubator at 37°C with 5% CO_2_. The growth medium was replaced every 2 days, and the cells were passaged every 4 days at a ratio of 1 to 4 upon 80% confluence. Human aortic vascular smooth muscle cells (VSMCs) (ScienCell, no. 6110) was routinely cultured in low-glucose DMEM. The culture method is similar to that of mouse primary VSMCs. Mdivi-1 (10 μmol/L; Enzo) was used to block mitochondrial fission.

### Senescence-associated β-galactosidase (SA-β-gal) staining

Cellular SA-β-gal activity was assayed using the Senescent Cells Staining Kit (Sigma, CS0030, USA) as previously described (Herbert *et al*, 2008). The SA-β-gal signals were analyzed using Image J software (NIH).

### Adenovirus expression vector and plasmid constructs

The expression plasmids of eIF5a and Mfn1were created by the placement of human eIF5a and Mfn1 cDNA into the pcDNA3.1 vector. The 5′ promoter regions of human eIF5a (−1500 to +1 bp) were amplified by PCR and cloned into the pGL3-basic vector (Promega) in order to generate the eIF5a promoter-reporter pGL3- eIF5a-luc. Truncated eIF5a luciferase reporters were generated by cloning the −1379, −674, −287 and −89 to +1 regions of the eIF5a promoter into pGL3. Adenoviruses encoding Klf5 (Ad-Klf5) and control (Ad-Ctl), Ad-shKlf5 and Ad-U6 were entrusted to Invitrogen. The VSMCs were washed and incubated in serum-free medium, and then were infected with the above adenovirus (5×10^9^ pfu/ml) for 36 h, followed by Ang II treatment.

### Small interfering RNA (siRNA) transfection

Small interfering RNAs (si-RNAs) targeting human eIF5a (si-eIF5a), Mfn1 (si-Mfn1) and Drp1 (si-Drp1) and negative control were designed and synthesized by GenePharma (Shanghai, China). The siRNAs were transiently transfected into VSMCs using the Lipofectamine 2000 reagent (Invitrogen). Twenty-hours following transfection, human VSMCs were treated with Ang II (100 ng/mL). Cells were then harvested and lysed for western blotting.

### Quantitative real-time PCR

Total RNA was extracted from VSMCs using TRizol Reagent and the cDNA was synthesized using the reverse transcriptase kit (Invitrogen). Real-time PCR was performed using SYBR Green RT-PCR Kit (Invitrogen). The relative mRNA expression was normalized to GAPDH and calculated using the 2^−ΔΔCt^ formula as previously described (Zhang *et al*, 2015). Sequence-specific primers used were presented at table 2.

**Table 2.**
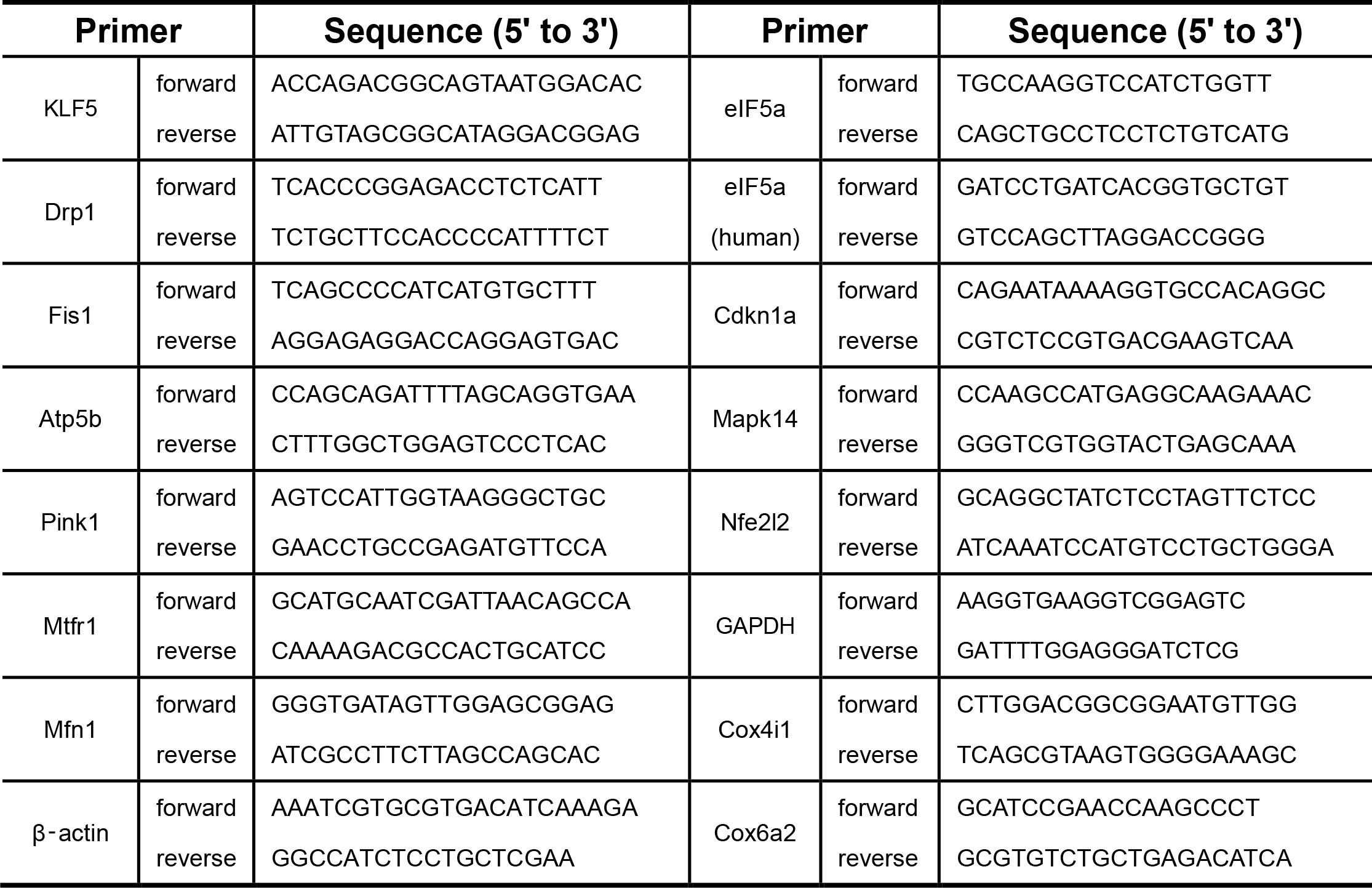
The primers for real-time PCR.

### Microarray analysis

Total RNA was extracted by using TRIzol® extraction method and a NucleoSpin RNA II kit (Macherey Nagel, Duren, Germany), from WT (n = 3) and smcKlf5^−/−^ (n = 3) mouse aortas at day 28. mRNA profile was assayed using the array hybridization as previously described. All gene level files were imported into Agilent GeneSpring GX software (version 12.1) for further analysis. Genes in 6 samples have values greater than or equal to lower cut-off: 100.0 (“All Targets Value”) were chosen for data analysis. GO Analysis was applied to determine the roles of these differentially expressed genes played in these biological GO terms.

### RNA-sequencing

RNA extraction of Ad-GFP- and Ad-Klf5-infected VSMCs was performed and sequenced on an Illumina HiSeq 2000. Illumina sequencing libraries were prepared according to the TruSeq RNA Sample Preparation Guide following the manufacturer’s instructions. Libraries were sequenced using 1 × 58 bp single-end reads, with two indexed samples per lane, yielding about 32.5 million reads per sample. After alignment of the sequencing reads to the human genome, counts for each gene were computed for each sample by use of the HTSeq software (version v0.5.3p3).

### Immunoblot analysis

Lysates from VSMCs were prepared and separated by SDS-PAGE, transferred to Immobilon-P membranes (Millipore), and incubated with specific antibodies. Western Lightning plus-ECL (PerkinElmer) was used for detection. Klf5 antibody (GTX103289, GeneTex) was from BD Biosciences. Antibodies of eIF5a (17069-1-AP), Fis1 (10956-1-AP), Mfn1 (13798-1-AP), Drp1 (12957-1-AP) and Mftr1 (bs-7632R) were from Proteintech (USA). β-actin (ab8226) were from Abcam (Cambridge, UK). Band intensities were quantified with the Image J software (NIH).

### Luciferase assay

Human embryonic kidney 293A cells were maintained as previously described (Li *et al*, 2010). 3×10^4^ VSMCs were seeded into each well of a 24-well plate and grown for 24 h prior to transfection with reporter plasmids and the control pTK-RL plasmid. VSMCs were transfected using Lipofectamine 2000 reagent (Invitrogen) according to the manufacturer’s instructions. Luciferase assays were performed after 24 h using a dual luciferase assay kit (Promega). Specific promoter activity was expressed as the relative ratio of firefly luciferase activity to Renilla luciferase activity. All promoter constructs were evaluated in a minimum of three separate wells per experiment.

### ChIP assay

VSMCs were cross-linked with 1% formaldehyde for 15 min, lysed as previously described, and then sonicated to an average size of 400–600 bp (Yu *et al*, 2011). The DNA fragments were immunoprecipitated overnight with anti-KLF5 or anti-mouse immunoglobulin G (IgG) (as a negative control). After the reversal of cross-linking, the eIF5a-flanking genomic region containing KLF5-binding sites was amplified by PCR; the forward and reverse primers used were as follows:

Primer1: Forward: 5′ ccagacagggcaaaggctcc 3′, Reverse: 5′ cggtgctacggctgcccctgc 3′;
Primer2: Forward: 5′ gcgaggtgagagcgggcagg 3′, Reverse: 5′ ggccccaaccgtcacccctc 3′;
Primer 3: Forward: 5′ -cgttgagacccaggcgtttc-3′, Reverse: 5′ ttccaaccagtcggcaatgc 3′.

### Oligonucleotide pull-down assay

Oligonucleotide pull-down assay was undertaken as described previously (Yu *et al*, 2011). Oligonucleotides containing the wild-type or mutant TCE site (site-1–3) sequence in the human eIF5a promoter with biotin added to their 5′-end were as follows:

site-1 (wild-type):

biotin-5’-gcgcggccacgtgaggtgggggagggggct-3’ (forward);
biotin-5’-agccccctcccccacctcacgtggccgcgc-3’ (reverse);
mutant site-1:

biotin-5’- gcgcggccacgtgagacaggggagggggct -3’(forward);
biotin-5’- agccccctcccctgtctcacgtggccgcgc -3’(reverse);
site-2 (wild-type):

biotin-5’- gccggccgggtgacgttggagcga -3’ (forward);
biotin-5’- tcgctccaacgtcacccggccggc -3’ (reverse);
mutant site-2:

biotin-5’- gccggccggacaacgttggagcga -3’ (forward);
biotin-5’- tcgctccaacgttgtccggccggc -3’ (reverse);
site-3 (wild-type):

biotin-5’-gctgtcgctcaaggggagggaagaagaggt-3’ (forward);
biotin-5’-acctcttcttccctccccttgagcgacagc-3’ (reverse)

### Co-immunoprecipitation assay

Co-immunoprecipitation was performed as described previously (Zhang *et al*, 2012). The cell lysates were immunoprecipitated with anti-eIF5a antibodies or anti-Mfn1, anti-Drp1, anti-Fis1 and anti-Mftr1 antibodies for 1 h at 4°C, followed by incubation with protein A-agarose overnight at 4°C. The precipitates were collected by centrifugation at 12,000 g for 1 min at 4°C, and washed 5 times with cold RIPA buffer before immunoblots using antibodies against eIF5a, Mfn1, Drp1, Fis1 and Mftr1.

### Proximity Ligation Assay

The experiment was performed as described previously (Ma *et al*, 2017). VSMCs were grown on cell culture inserts and incubated with 4% paraformaldehyde for 10 minutes. Proximity ligation assay was performed using the Rabbit PLUS and Mouse MINUS Duolink in situ proximity ligation assay kits with anti-eIF5a (mouse) and anti-Mfn1 (rabbit; OLINK Bioscience, Uppsala, Sweden) according to the manufacturer’s protocol. Subsequently, slides were dehydrated, air-dried, and embedded in 4’,6-diamidino-2-phenylindole-containing mounting medium.

### Cell immunofluorescence

Immunofluorescence staining was performed as described previously (Ma *et al*, 2017). Cells were permeabilized with 0.1% Triton X-100 in PBS and then blocked with 0.1% Triton X-100 and 5% BSA in PBS for 1 h, washed, and incubated overnight with the primary antibodies anti-eIF5a (1:50, sc-390062, Santa), anti-Drp1 (1:100, 12957-1-AP, Proteintech), anti-Mfn1 (1:100, 13798-1-AP, Proteintech), anti-Fis (1:100, 10956-1-AP, Proteintech), and anti-Mftr1 (1:50, bs-7632R, Bioss) at 4°C. Antibodies conjugated to Alexa Fluor 405, 488 and 568 were used as secondary antibodies. Mitochondrial was visualized with Mitotracker green and nuclei were stained with 4′-6-diamidino-2-phenylindole (DAPI). Images were captured by confocal microscopy (DM6000 CFS, Leica) and processed by LAS AF software. All images of primary cells were taken with 63×NA 0.75 objectives. One hundred cells were counted for each assay.

### Measurement of total and mitochondrial ROS levels

The ROS measurement was performed as previously described (Luo *et al*, 2017). ROS level was evaluated by analyzing the fluorescence intensity that resulted from dihydroethidium (DHE) (Invitrogen) and mitoSOX (Invitrogen) staining. In brief, frozen mouse aortas were cut into 4 μm sections. Serial aorta sections were stained with 5 μM DHE or mitoSOX at 37°C for 30 min and then measured by fluorescence microscopy. The treated VSMCs were loaded with 5 μM DHE or mitoSOX at 37°C for 30 min and then measured by fluorescence microscopy.

### ATP assay

ATP levels were determined using the Adenosine 5′-triphosphate (ATP) Bioluminescent Assay Kit (Sigma) following the manufacturer’s instructions. Luminescence was recorded using a luminometer (Molecular Probes).

### TUNEL assay

A terminal deoxynucleotidyl-transferase-mediated dUTP nick end-labeling (TUNEL) assay kit (FITC, ab66108) was used to identify apoptosis cells in aorta tissues following the manufacturer’s instructions. The FITC-labeled TUNEL-positive cells in aorta tissues were imaged under a fluorescent microscopy by using 488 nm excitation and 530 nm emission. Apoptosis of human VSMCs was detected by flow cytometry. In brief, the cells were harvested by trypsinization, and washed twice with cold PBS. Next, the cells were centrifuged and resuspended in the binding buffer at a density of 1.0 × 10^6^ cells/mL. The cells were incubated with 5 μL of annexin V-FITC and 5 μL of PI for 15 min at room temperature in the dark. All samples were subject to flow cytometry (FC500 MPL Beckman) and analyzed with the FlowJo software.

### Transmission electron microscopy

Samples were processed following standard protocol, including dehydration, embedding, and sectioning, and then examined and captured under a Hitachi JEM-1400 transmission electron microscope (Hitachi, Tokyo, Japan). The number of mitochondrial was quantified by counting the total number of mitochondria per 100 μm^2^ of cell surface. Mitochondrial mean diameter was calculated as the average diameter per mitochondrion, 200 mitochondria per group were randomly selected. Mitochondrial major/minor axes length was quantified using NIH Image J software.

### Statistical analysis

For each experiment, three or more fields of view were taken as fluorescent images for each group. Fluorescent intensity in each image was semi-quantitated with ImageJ and averaged. Results were displayed as mean fluorescence intensity. All data are displayed as representative or results from multiple independent experiments. Data comparisons were performed with one-way analysis of variance test. Two-sided P values of <0.05 were considered significant and denoted with 1, 2, or 3 asterisks when lower than 0.05, 0.01, or 0.001, respectively. No randomization or blinding was used.

**Expanded View** for this article is available online.

## Acknowledgements

This work was supported by grants from the National Natural Science Foundation of China (No. 31671182, No. 31871152 and No. 81700416).

## Author contributions

Conception and design: DM. and J-KW; Acquisition of data (including provided animals, provided facilities, etc.): BZ, H-LL, Y-BZ and XL; Technical assistance: TS; Writing the manuscript: DM and J-KW; Study supervision: J-KW.

## References

Abuarab N, Munsey TS, Jiang LH, Li J, Sivaprasadarao A (2017) High glucose–induced ROS activates TRPM2 to trigger lysosomal membrane permeabilization and Zn2^+^-mediated mitochondrial fission. Sci Signal 4161: 1–12

Anderson GR, Wardell SE, Cakir M, Yip C, Ahn YR, Ali M, Yllanes AP, Chao CA, McDonnell DP, Wood KC (2018) Dysregulation of mitochondrial dynamics proteins are a targetable feature of human tumors. Nat Commun 26: 1677–1689

Aoki T, Nishimura M, Kataoka H, Ishibashi R, Nozaki K, Hashimoto N (2009) Reactive oxygen species modulate growth of cerebral aneurysms: a study using the free radical scavenger edaravone and p47phox(-/-) mice. Lab Invest 89: 730–741

Alcolea PJ, Alonso A, García-Tabares F, Toraño A, Larraga V (2014) An Insight into the proteome of Crithidia fasciculata choanomastigotes as a comparative approach to axenic growth, peanut lectin agglutination and differentiation of Leishmania spp. promastigotes. PLoS One 9: e113837

Andrés V (2014) Vitamin D puts the brakes on angiotensin II-induced oxidative stress and vascular smooth muscle cell senescence. Atherosclerosis 236: 444–447

Benne R, Brown-Luedi ML, Hershey JW (1978) Purification and characterization of protein synthesis initiation factors eIF-1, eIF-4C, eIF-4D, and eIF-5 from rabbit reticulocytes. J Biol Chem 253: 3070–3077

Bordt EA, Clerc P, Roelofs BA, Saladino AJ, Tretter L, Adam-Vizi V, Cherok E, Khalil A, Yadava N, Ge SX, et al (2017) The Putative Drp1 Inhibitor mdivi-1 Is a Reversible Mitochondrial Complex I Inhibitor that Modulates Reactive Oxygen Species. Dev Cell 40: 583–594.e6

Chi C, Li DJ, Jiang YJ, Tong J, Fu H, Wu YH, Shen FM (2019) Vascular smooth muscle cell senescence and age-related diseases: State of the art. Biochim Biophys Acta Mol Basis Dis 1865: 1810–1821

Caja S and Enríquez JA (2017) Mitochondria in endothelial cells: Sensors and integrators of environmental cues. Redox Biol 12: 821–827

Drosatos K, Pollak NM, Pol CJ, Ntziachristos P, Willecke F, Valenti MC, Trent CM, Hu Y, Guo S, Aifantis I, et al (2016) Cardiac Myocyte KLF5 Regulates Ppara Expression and Cardiac Function. Circ Res 118: 241–253

Dzau VJ (2001) Theodore Cooper Lecture: Tissue angiotensin and pathobiology of vascular disease: a unifying hypothesis. Hypertension 37: 1047–1052

Galloway CA, Lee H, Nejjar S, Jhun BS, Yu T, Hsu W, Yoon Y (2012) Transgenic control of mitochondrial fission induces mitochondrial uncoupling and relieves diabetic oxidative stress. Diabetes 61: 2093–2104

He M, Han M, Zheng B, Shu YN, Wen JK (2009) Angiotensin II stimulates KLF5 phosphorylation and its interaction with c-Jun leading to suppression of p21 expression in vascular smooth muscle cells. J Biochem 146: 683–691

Herbert KE, Mistry Y, Hastings R, Poolman T, Niklason L, Williams B (2008) Angiotensin II-mediated oxidative DNA damage accelerates cellular senescence in cultured human vascular smooth muscle cells via telomere-dependent and independent pathways. Circ Res 102: 201–208

Hernandez-Segura A, Nehme J, Demaria M (2018) Hallmarks of Cellular Senescence. Trends Cell Biol 28: 436–453

Hoque A, Sivakumaran P, Bond ST, Ling NXY, Kong AM, Scott JW, Bandara N, Hernández D, Liu GS, Wong RCB, et al (2018) Mitochondrial fission protein Drp1 inhibition promotes cardiac mesodermal differentiation of human pluripotent stem cells. Cell Death Discov 4: 39–41

Kim S, Kim C, Park S (2017) Mdivi-1 Protects Adult Rat Hippocampal Neural Stem Cells against Palmitate-Induced Oxidative Stress and Apoptosis. Int J Mol Sci 18: pii: E1947

Kunieda T, Minamino T, Nishi J, Tateno K, Oyama T, Katsuno T, Miyauchi H, Orimo M, Okada S, Takamura M, et al (2006) Angiotensin II induces premature senescence of vascular smooth muscle cells and accelerates the development of atherosclerosis via a p21-dependent pathway. Circulation 114: 953–960

Lacolley P, Regnault V, Segers P, Laurent S (2017) Vascular Smooth Muscle Cells and Arterial Stiffening: Relevance in Development, Aging, and Disease. Physiol Rev 97:1555–1617

Lakatta EG and Levy D (2003) Arterial and cardiac aging: major shareholders in cardiovascular disease enterprises: Part II: the aging heart in health: links to heart disease. Circulation 107: 346–354

Li DJ, Huang F, Ni M, Fu H, Zhang LS, Shen FM (2016) α7 Nicotinic Acetylcholine Receptor Relieves Angiotensin II-Induced Senescence in Vascular Smooth Muscle Cells by Raising Nicotinamide Adenine Dinucleotide-Dependent SIRT1 Activity. Arterioscler Thromb Vasc Biol 36: 1566–1576

Li HX, Han M, Bernier M, Zheng B, Sun SG, Su M, Zhang R, Fu JR, Wen JK (2010) Krüppel-like factor 4 promotes differentiation by transforming growth factor-beta receptor-mediated Smad and p38 MAPK signaling in vascular smooth muscle cells. J Biol Chem 285: 17846–17856

Liu Y, Wen JK, Dong LH, Zheng B, Han M (2010) Krüppel-like factor (KLF) 5 mediates cyclin D1 expression and cell proliferation via interaction with c-Jun in Ang II-induced VSMCs. Acta Pharmacol Sin 31: 10–18

Lubas M, Harder LM, Kumsta C, Tiessen I, Hansen M, Andersen JS, Lund AH, Frankel LB (2018) eIF5A is required for autophagy by mediating ATG3 translation. EMBO Rep 19: e46072

Luchessi AD, Cambiaghi TD, Alves AS, Parreiras-E-Silva LT, Britto LR, Costa-Neto CM, Curi R (2008) Insights on eukaryotic translation initiation factor 5A (eIF5A) in the brain and aging. Brain Res 1228: 6–13

Luo YX, Tang X, An XZ, Xie XM, Chen XF, Zhao X, Hao DL, Chen HZ, Liu DP (2017) Sirt4 accelerates Ang II-induced pathological cardiac hypertrophy by inhibiting manganese superoxide dismutase activity. Eur Heart J 38: 1389–1398

Ma D, Zheng B, Suzuki T, Zhang R, Jiang C, Bai D, Yin W, Yang Z, Zhang X, Hou L, Zhan H, Wen JK (2017) Inhibition of KLF5-Myo9b-RhoA Pathway-Mediated Podosome Formation in Macrophages Ameliorates Abdominal Aortic Aneurysm. Circ Res 120: 799–815

Makino A, Scott BT, Dillmann WH (2010) Mitochondrial fragmentation and superoxide anion production in coronary endothelial cells from a mouse model of type 1 diabetes. Diabetologia 53: 1783–1794

Miao SB, Xie XL, Yin YJ, Zhao LL, Zhang F, Shu YN, Chen R, Chen P, Dong LH, Lin YL (2017) Accumulation of Smooth Muscle 22α Protein Accelerates Senescence of Vascular Smooth Muscle Cells via Stabilization of p53 In Vitro and In Vivo. Arterioscler Thromb Vasc Biol 37: 1849–1859

Miyake T, Pradeep S, Wu SY, Rupaimoole R, Zand B, Wen Y, Gharpure KM, Nagaraja AS, Hu W, Cho MS, et al (2015) XPO1/CRM1 Inhibition Causes Antitumor Effects by Mitochondrial Accumulation of eIF5A. Clin Cancer Res 21: 3286–3297

Muñoz-Espín D, Serrano M (2014) Cellular senescence: from physiology to pathology. Nat Rev Mol Cell Biol 15: 482–496

Murphy MP (2009) How mitochondria produce reactive oxygen species. Biochem J 417: 1–13

Nguyen S, Leija C, Kinch L, Regmi S, Li Q, Grishin NV, Phillips MA (2015) Deoxyhypusine Modification of Eukaryotic Translation Initiation Factor 5A (eIF5A) Is Essential for Trypanosoma brucei Growth and for Expression of Polyprolyl-containing Proteins. J Biol Chem 290: 19987–19998

Nordon IM, Hinchliffe RJ, Loftus IM, Thompson MM (2011) Pathophysiology and epidemiology of abdominal aortic aneurysms. Nat Rev Cardiol 8: 92–102

Patel PH, Costa-Mattioli M, Schulze KL, Bellen HJ (2009) The Drosophila deoxyhypusine hydroxylase homologue nero and its target eIF5A are required for cell growth and the regulation of autophagy. J Cell Biol 185: 1181–1194

Pereira KD, Tamborlin L, Meneguello L, de Proença AR, Almeida IC, Lourenço RF, Luchessi AD (2016) Alternative Start Codon Connects eIF5A to Mitochondria. J Cell Physiol 231: 2682–2689

Przybylska D, Janiszewska D, Goździk A, Bielak-Zmijewska A, Sunderland P, Sikora E, Mosieniak G (2016) NOX4 downregulation leads to senescence of human vascular smooth muscle cells. Oncotarget 7: 66429–66443

Puleston DJ, Buck MD, Klein Geltink RI, Kyle RL, Caputa G, O’Sullivan D, Cameron AM, Castoldi A, Musa Y, Kabat AM, et al (2019) Polyamines and eIF5A Hypusination Modulate Mitochondrial Respiration and Macrophage Activation. Cell Metab 30: 30243–30258

Quintana RA and Taylor WR (2019) Cellular Mechanisms of Aortic Aneurysm Formation. Circ Res 124: 607–618

Raffort J, Lareyre F, Clément M, Hassen-Khodja R, Chinetti G, Mallat Z (2017) Monocytes and macrophages in abdominal aortic aneurysm. Nat Rev Cardiol 14: 457–471

Salmon M, Johnston WF, Woo A, Pope NH, Su G, Upchurch GR Jr, Owens GK, Ailawadi G (2013) KLF4 Regulates Abdominal Aortic Aneurysm Morphology and Deletion Attenuates Aneurysm Formation. Circulation 128: S163–S174

Seko Y, Fujimura T, Yao T, Taka H, Mineki R, Okumura K, Murayama K (2015) Secreted tyrosine sulfated-eIF5A mediates oxidative stress-induced apoptosis. Sci Rep 5: 13737–13751

Serasinghe MN and Chipuk JE (2017) Mitochondrial Fission in Human Diseases. Handb Exp Pharmacol 240:159–188

Shenouda SM, Widlansky ME, Chen K, Xu G, Holbrook M, Tabit CE, Hamburg NM, Frame AA, Caiano TL, Kluge MA, et al (2011) Altered mitochondrial dynamics contributes to endothelial dysfunction in diabetes mellitus. Circulation 124: 444–453

Shindo T, Manabe I, Fukushima Y, Tobe K, Aizawa K, Miyamoto S, Kawai-Kowase K, Moriyama N, Imai Y, Kawakami H, et al (2002) Krüppel-like zinc-finger transcription factor KLF5/BTEB2 is a target for angiotensin II signaling and an essential regulator of cardiovascular remodeling. Nat Med 8: 856–863

Shi Y, Fan S, Wang D, Huyan T, Chen J, Chen J, Su J, Li X, Wang Z, Xie S, et al (2018) FOXO1 inhibition potentiates endothelial angiogenic functions in diabetes via suppression of ROCK1/Drp1-mediated mitochondrial fission. Biochim Biophys Acta Mol Basis Dis 1864, 2481–2494

Suzuki T, Aizawa K, Matsumura T, Nagai R (2005) Vascular implications of the Krüppel-like family of transcription factors. Arterioscler Thromb Vasc Biol 25:1135–1141

Suzuki T, Sawaki D, Aizawa K, Munemasa Y, Matsumura T, Ishida J, Nagai R (2009) Kruppel-like factor 5 shows proliferation-specific roles in vascular remodeling, direct stimulation of cell growth, and inhibition of apoptosis. J Biol Chem 284: 9549–9557

Tezze C, Romanello V, Desbats MA, Fadini GP, Albiero M, Favaro G, Ciciliot S, Soriano ME, Morbidoni V, Cerqua C, et al (2017) Age-Associated Loss of OPA1 in Muscle Impacts Muscle Mass, Metabolic Homeostasis, Systemic Inflammation, and Epithelial Senescence. Cell Metab 25: 1374–1389.e6

Valcheva P, Cardus A, Panizo S, Parisi E, Bozic M, Lopez Novoa JM, Dusso A, Fernández E, Valdivielso JM (2014) Lack of vitamin D receptor causes stress-induced premature senescence in vascular smooth muscle cells through enhanced local angiotensin-II signals. Atherosclerosis 235: 247–255

Vásquez-Trincado C, García-Carvajal I, Pennanen C, Parra V, Hill JA, Rothermel BA, Lavandero S (2016) Mitochondrial dynamics, mitophagy and cardiovascular disease. J Physiol 594: 509–525

Yang YC, Tsai CY, Chen CL, Kuo CH, Hou CW, Cheng SY, Aneja R, Huang CY, Kuo WW (2018) Pkcδ Activation is Involved in ROS-Mediated Mitochondrial Dysfunction and Apoptosis in Cardiomyocytes Exposed to Advanced Glycation End Products (Ages). Aging Dis 9: 647–663

Yu K, Zheng B, Han M, Wen JK (2011) ATRA activates and PDGF-BB represses the SM22α promoter through KLF4 binding to, or dissociating from, its cis-DNA elements. Cardiovasc Res 90: 464–474

Zhang XH, Zheng B, Gu C, Fu JR, Wen JK (2012) TGF-β1 downregulates AT1 receptor expression via PKC-δ-mediated Sp1 dissociation from KLF4 and Smad-mediated PPAR-γ association with KLF4. Arterioscler Thromb Vasc Biol 32: 1015–1023

Zhang XH, Zheng B, Yang Z, He M, Yue LY, Zhang RN, Zhang M, Zhang W, Zhang X, Wen JK (2015) TMEM16A and myocardin form a positive feedback loop that is disrupted by KLF5 during Ang II-induced vascular remodeling. Hypertension 66: 412–421

Zhao J, Liu T, Jin S, Wang X, Qu M, Uhlén P, Tomilin N, Shupliakov O, Lendahl U, Nistér M (2011) Human MIEF1 recruits Drp1 to mitochondrial outer membranes and promotes mitochondrial fusion rather than fission. EMBO J 30: 2762–2778

Zheng B, Han M, Shu YN, Li YJ, Miao SB, Zhang XH, Shi HJ, Zhang T, Wen JK (2011) HDAC2 phosphorylation-dependent Klf5 deacetylation and RARα acetylation induced by RAR agonist switch the transcription regulatory programs of p21 in VSMCs. Cell Res 21: 1487–1508

Zheng B, Yin WN, Suzuki T, Zhang XH, Zhang Y, Song LL, Jin LS, Zhan H, Zhang H, Li JS, Wen JK (2017) Exosome-Mediated miR-155 Transfer from Smooth Muscle Cells to Endothelial Cells Induces Endothelial Injury and Promotes Atherosclerosis. Mol Ther 25: 1279–1294

